# scHumanNet: a single-cell network analysis platform for the study of cell-type specificity of disease genes

**DOI:** 10.1101/2022.06.20.496836

**Authors:** Junha Cha, Jiwon Yu, Jae-Won Cho, Martin Hemberg, Insuk Lee

## Abstract

A major challenge in single-cell biology is identifying cell-type-specific gene functions, which may substantially improve precision medicine. Differential expression analysis of genes is a popular, yet insufficient approach, and complementary methods that associate function with cell type are required. Here, we describe scHumanNet (https://github.com/netbiolab/scHumanNet), a single-cell network analysis platform for resolving cellular heterogeneity across gene functions in humans. Based on cell-type-specific networks (CSNs) constructed under the guidance of the HumanNet reference interactome, scHumanNet displayed higher functional relevance to the cellular context than CSNs built by other methods on single-cell transcriptome data. Cellular deconvolution of gene signatures based on network compactness across cell types revealed breast cancer prognostic markers associated with T cells. scHumanNet could also prioritize genes associated with particular cell types using CSN centrality and identified the differential hubness of CSNs between disease and healthy conditions. We demonstrated the usefulness of scHumanNet by uncovering T-cell-specific functional effects of *GITR*, a prognostic gene for breast cancer, and functional defects in autism spectrum disorder genes specific for inhibitory neurons. These results suggest that scHumanNet will advance our understanding of cell-type specificity across human disease genes.

## Introduction

Genes do not act in isolation, because the proteins they encode interact with each other and with other molecules. From the perspective of network biology, molecular interactions determine the function of each cell type (1). However, cell-type-specific molecular interactions are difficult to identify and interpret due to context dependency. The advent of single-cell RNA sequencing (scRNA-seq) has enabled the characterization of distinct cell types from complex tissues, as well as the determination of their interactions within mixed-cell populations (2).

A major difficulty in cell-type-specific network (CSN) inference from single-cell transcriptome data is the lack of a gold standard for cell-type-specific gene interactions. Accordingly, researchers often use simulated synthetic networks (3). An evaluation using reference protein-protein interactions showed that most methods for network inference, including those developed for bulk RNA-seq data and scRNA-seq, were not capable of reconstructing accurate networks of gene interactions from scRNA-seq data (4). This poor performance is likely due to elevated sparsity (5) and spurious technical variation (6) among scRNA-seq data. To overcome this problem, an accurate network modeling method that uses scRNA-seq data to study cell-type-specific gene functions should be developed.

Two approaches to network construction using single-cell transcriptome data exist: reference-free and reference-guided inference. The former, which is more popular, enables the discovery of gene interactions directly from single-cell transcriptome data, but it suffers from a generally high false-positive rate (4, 7). In contrast, the reference-guided approach builds a network by filtering the reference interactome for a given transcriptome of context-associated single cells. Filtered interactions are highly likely to exist in a given cell type.

Here, we describe scHumanNet, a computational platform for the reference-guided construction of CSNs using single-cell transcriptome data. As the reference interactome, we used HumanNet (8), one of the best-performing human gene networks for disease gene predictions. We utilized a modified version of the SCINET algorithm (9). Along with CSN construction, scHumanNet provides several analytical tools to aid the study of cell-type-specific effects of disease genes. Through network centrality analysis, we found that scHumanNet outperformed other single-cell network inference methods in retrieving cell-type-specific genes, suggesting it was suitable for the study of gene cell-type specificity. We demonstrated that genes relevant to the same cell type showed higher within-group connectivity (i.e., compactness) within the network. Utilizing network compactness across CSNs, we deconvolved breast cancer prognostic signatures into cell types and identified those associated with immune cells rather than cancer cells. We also found that the prognostic value of a known signature gene, *GITR*, was linked to T cells owing to its T-cell-specific centrality. Furthermore, we developed a statistical framework for differential centrality analysis that revealed cell-type-specific functional defects in disease genes. Applying this analytical framework to brain scRNA-seq data from autism studies, we found elevated dysregulation of the interaction networks in inhibitory and excitatory neurons of disease condition.

## Materials and Methods

### Single-cell transcriptome data for network construction

To construct CSNs, we used scRNA-seq data generated from biopsy samples of breast, lung, colorectal, and ovarian cancers with cell type annotations obtained from Qian *et al*. (10). For pan-cancer comparative network analysis, we focused on five major cell types in the tumor microenvironment: T cells, B cells, myeloid cells, cancer-associated fibroblasts (CAFs), and endothelial cells (ECs). For the study of autism spectrum disorder (ASD), we constructed CSNs for cell types found in the brain using scRNA-seq data obtained from Velmeshev *et al.* (11). The pre-annotated cell types were merged with more granular representations to include ECs, oligodendrocytes, astrocytes (AST-FB and AST-PP), microglia, inhibitory cells (IN-PV, IN-SST, IN-SV2C, and IN-VIP), excitatory cells (L2/3, L4, L5/6, and L5/6-CC), and others (Neu-mat, Neu-NRGN-I, and Neu-NRGN-II).

### Reference-free CSN construction with single-cell transcriptome data

For network construction, we only considered protein-coding genes defined by the consensus coding sequence (CCDS) database. We built four variants of the co-expression networks, each of them based on each scRNA-seq dataset. In the first co-expression network, we calculated Pearson correlation coefficients (*PCC*) between gene pairs using a count matrix of single-cell transcriptome data, which was log-normalized by the *NormalizeData()* function of the Seurat package. Only links with *PCC* > 0.8 were retained for the rawPCC network. The second type of co-expression network was based on the de-noised count matrix from MetaCell (12). To calculate the *PCC* between gene pairs, we used metacells generated with unified threshold and parameters. We discarded cells with fewer than 500 UMIs and used the parameters K = 30 and alpha = 2 for *mcell_mc_from_coclust_balanced()*. The third type of co-expression network was based on a count matrix with imputation of dropouts by SAVER (13) and exclusion of genes with >99% zero values. The last type of co-expression network was based on data transformation using bigSCale2 (14). The *recursive* method was used for the clustering parameter of *compute.network()*, and the PCC was calculated using the transformed Z-score matrix.

The accuracy of co-expression networks based on metacells, SAVER, and bigSCale2 was evaluated using a Bayesian statistical framework and log likelihood score (*LLS*) (15). In this scheme, gold standard gene pairs were used to evaluate the likelihood of data-driven gene pairs such as co-expression links. In brief, for the prioritized gene pairs inferred from the given data (*D*), we calculated *LLS* for every 1,000 links sorted by the data intrinsic score using the following equation:

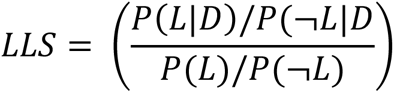

where P(*L*|*D*) and P(¬*L*|*D*) account for the probability of positive and negative gold standard gene pairs in a given dataset, respectively, and P(*L*) and P(¬*L*) represent the probability of gold standard positive and negative gene pairs, respectively. We used a set of 260,962 gold standard positive gene pairs obtained from HumanNet (8). A set of gold standard negative gene pairs was inferred as being composed of all links not included among gold standard positives.

For the construction of CSNs using GRNboost2 (16), 2,416 transcription factors (TFs) gathered from previous publications (17, 18) were used as input, and the top 0.1% of links were retained for the final networks.

### Reference-guided CSN construction with single-cell transcriptome data

scHumanNet was developed by modifying the SCINET framework (9) which utilizes imputation, transformation, and normalization of scRNA-seq data. Single-cell gene expression data were pre-processed using the ACTIONet package (19). By identifying the archetypes within the scRNA-seq dataset, ACTIONet learns the dominant transcriptional patterns representative of cell types and states. This approach produces a transformed gene activity score matrix, which is the basis for inferring gene-pair interactions. For each gene pair from the gene score activity matrix, a minimum activity score threshold is applied to assess the strength of the interactions in a group of cells. If each gene in the examined interaction passes the threshold determined by the transformed cell type activity score and a link exists in the reference interactome, it is deemed cell-type-specific and retained in the resultant CSN. Although the SCINET package provides edge weights based on the aggregated *p*-value of a likelihood score, we used the *LLS* from our reference interactome, HumanNet, as the edge weight. We measured the network centrality of each gene based on the sum of *LLSs* to all its neighbors. Because the human interactome is biased towards the ribosome complex (20), we excluded ribosomal proteins from the final candidate hub genes.

### Significance test for network hubness

The statistical significance of hub genes was calculated using the *FindAllHub()* function in the scHumanNet package. For each network, random networks were generated by swapping edges with equal probability, and the centrality scores of all genes were collected. This process was iterated until at least 10,000 centrality scores were gathered, which were then used to generate a null distribution. By default, Benjamini–Hochberg correction was applied for each *p*-value, and hub genes with false discovery rate (FDR) < 0.05 were selected for each CSN.

### Predicting cell-type-specific genes for B and T cells

To test the cell-type relevance of genes, we compiled T- and B-cell-associated genes from the Gene Ontology (GO) database. We used reliable annotations by considering only evidence based on traceable author statement (TAS), inferred from direct assay (IDA), inferred from mutant phenotype (IMP), or inferred from genetic interaction (IGI). By selecting GO term descriptions that contained either ‘T cell’ or ‘B cell’, we obtained 289 genes associated with T cells and 89 with B cells. We conducted a similar compilation for other cell types but could not obtain enough associated genes for statistical testing. We identified differentially expressed genes (DEGs) using the function *FindAllMarkers()* from the Seurat v3.2.3 package with default parameters ‘wilcox’ for *test.use*, ‘0.25’ for *logfc.threshold*, and ‘0.1’ for *min.pct*. We selected protein-coding DEGs with positive log-fold changes for B or T cells (*q*-value < 0.05) as cell-type-specific genes. Finally, we measured the weighted degree centrality of genes using the sum of edge scores for other network construction methods: *PCC* (rawPCC, MetaCell, SAVER, and bigSCale2), importance score (GRNboost2), and weighted score (SCINET). Only significant DEGs and hub genes were used to compare cell-type relevance.

### Predicting cell-type-specific TFs

TFs specific for B and T cells were obtained from the TF-Marker database (21) and subsequently filtered using the TRRUST database (22), resulting in 42 T-cell-associated TFs and 14 B-cell-associated TFs. The top 100 hub genes identified by scHumanNet were extracted from each cell type and filtered using the 2,416 TFs collated from previous publications (17, 18). The top 100 DEGs based on log-fold change values were selected and filtered using the same TF gene list. Hypergeometric tests were performed with all genes in HumanNet (18, 593) as the gene space.

### Compactness analysis of gene sets to identify relevant cell types

We implemented the *Connectivity()* function in scHumanNet to evaluate network compactness of a group of genes. Briefly, 10,000 random gene sets with the same number of genes as the test gene set were selected to generate a null model. To preserve the network topological properties for the random gene sets, we used rejection sampling, whereby we selected a gene with ±20% degree of connectivity for each real gene when permuting. Significance was measured using the rank of observed within-group connectivity in the null distribution. Genes that exert their function in a specific cell type tend to be connected to each other in a network specific to the cell type. The degree of compactness was measured using the significance of within-group connectivity. We performed compactness analysis for a set of immune checkpoint molecule (ICM) genes (23) and 33 breast cancer prognostic signature gene sets collected from Huang *et al.* (24). For the GGI97 signature, only 76 out of 97 genes were evaluated in this study because the others had been either discontinued or deprecated in the NCBI gene database (**Supplemental Table 1**). Their relevance to the cell cycle was assessed using manual curation and accepted databases. Genes that were included in ‘Cell Cycle’ of KEGG 2021, ‘G2-M Checkpoint’ of MSigDB 2020, and ‘Cell Cycle Homo sapiens’ of Reactome DB 2016 were considered cell cycle-related. Other genes were curated manually, and those that included ‘DNA replication’ and ‘mitotic spindle’ were also included. Of the 76 signature genes, 24 were detected in the breast cancer T-cell network, and their functional connectivity was assessed through *Connectivity()* with default parameters.

### Survival analysis on The Cancer Genome Atlas (TCGA) breast cancer samples

Only direct neighbors of the *GITR* gene in the T-cell network for breast cancer were considered connected to GGI97 signatures. TCGA data were downloaded through the GDC portal using the *TCGAbiolinks* R package. HTseq counts were preprocessed using *TCGAanlayze_Preprocessing()*, with ‘0.6’ as the *cor.cut* parameter. The data were subsequently normalized using *TCGAanalyze_Normalization()*. The preprocessed count data were normalized with sample-specific size factors calculated using DESeq2. To identify genes indicative of good patient outcomes, we considered 23,192 genes from TCGA-derived expression matrix, of which 1,078 BRCA samples were separated based on the top 30th and bottom 30th percentile of test gene expression. *P*-values from the Kaplan–Meier log test were corrected using the Benjamini– Hochberg method, yielding 236 genes with FDR < 0.05, which were regarded as predictive of good prognosis. For survival analysis, samples were separated into high and low groups based on median *GITR* expression. The correlation between *GITR* expression for each bulk sample and the composition of T cells was calculated using the geometric mean of *CD3D*, *CD3E*, and *CD3G*. The survival group was divided into high and low groups based on the median of either single gene expression or geometric mean expression of the gene set. Network visualization of the breast cancer T-cell scHumanNet was performed using the *igraph* R package.

### Differential centrality analysis for ASD genes

For each CSN, the degree of centrality was assessed based on the sum of edge weights (*LLS*). Because network size affects the degree of centrality score, we used percentile ranks, whereby the most central gene had a value of 1 and the least central one had a value of 0. We assigned a value of 0 to genes that were not included in at least one of the networks. For each gene, we calculated the differential percentile rank of centrality (diffPR) by subtracting the percentile rank in the control network from the percentile rank in the disease network.

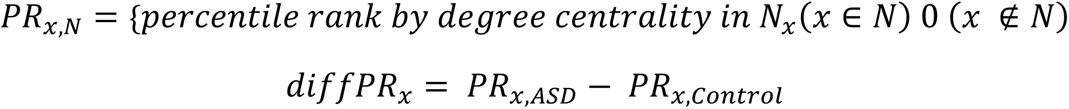

where *x* represents a gene and *N* represents a disease or control network for a given cell type.

The percentile rank was calculated using the *dplyr* package *percent_rank().* The *diffPR* for each gene ranged from -1 to 1, with positive values indicating higher connectivity in the disease network.

For a significance test of differential centrality, we used the *FindDiffHub()* and *TopDiffHub()* functions in scHumanNet. Briefly, *FindDiffHub()* finds a distribution of null *diffPR* values for every gene by random permutation of the control network to measure the significance of the observed differential centrality. Random sampling of *diffPR* values continues until one million random values accumulate. Benjamini–Hochberg correction was applied to calculate the FDR. For *TopDiffHub(),* the *diffPR* of the genes was assessed and filtered for non-zero values. By default, genes within the top 5% of *diffPR* values were selected as differential hub genes. To define lost and gained hub genes in the disease network, 0.7 was set as the threshold. Accordingly, control hub genes with a percentile rank > 0.7 were assessed for their *diffPR* distribution. We observed a clear bimodal pattern dividing the genes around a specific *diffPR* value. Genes with *diffPR* of the same threshold or above (absolute value) were considered as hub genes and were characterized by large changes between healthy controls and disease CSNs. Functional enrichment analysis was performed using the *enrichR* package (25) with pathway terms derived from five pathway databases: Elsevier Pathway Collection (as of March 2022), BioPlanet v.1.0, Reactome 2016, GO Biological Process (GOBP) (as of March 2022), and GO Molecular Function (GOMF) (as of March 2022).

## Results

### scHumanNet effectively retrieves genes specific for each intratumoral cell type

To evaluate whether CSNs obtained by scHumanNet (**Figure 1A**) were more suitable than those generated by other inference methods for the study of cell-type-specific gene functions, we compared various reference-free and reference-guided approaches. Using published breast cancer scRNA-seq data (10), we constructed networks for T cells, B cells, myeloid cells, ECs, CAFs, and cancer cells using five reference-free methods, including rawPCC, MetaCell (12), SAVER (13), GRNboost2 (16), and bigSCale2 (14), as well as one reference-guided method, SCINET (9) based on PCNet (26). Network size across cell types and network inference methods varied widely (**Supplemental Table 2**).

**Figure 1.**
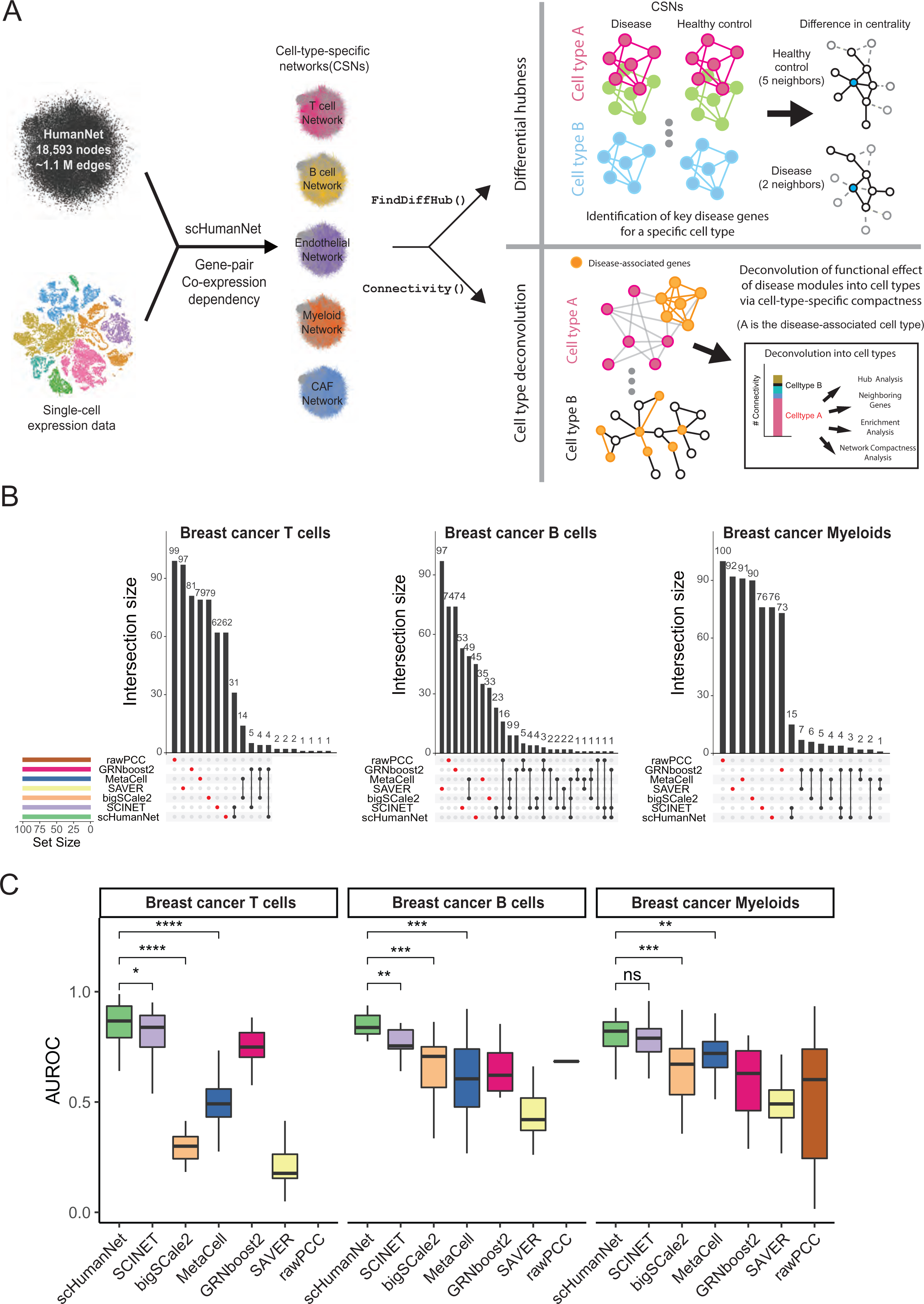
Comparison of cell-type-specific networks generated by scHumanNet and other methods. A. Overview of the scHumanNet platform and downstream analysis scheme used in this study **B.** Upset plots of the top 100 hub genes in breast cancer networks specific for T cells, B cells, and myeloid cells constructed using seven different network inference methods. **C.** Area under the receiver operating characteristic curve (AUROC) used to assess retrieval of cell-type-specific genes derived from the Azimuth cell type database by centrality in T, B, and myeloid cell-specific networks of breast cancer. \**p* < 0.05, \*\**p* < 0.01, \*\*\**p* < 0.001.

The functional importance of network nodes is measured by their centrality. Cell-type-specific genes presumably play important roles in the corresponding cell types. Therefore, we expected that genes with high centrality values in each CSN were enriched for cell-type-specific genes. Using weighted degree centrality based on the edge scores of each network, we compared the top 100 genes from each network. In this way, we could disregard differences in network size. Interestingly, no overlap was observed between any of the six network construction methods when assessing the top 100 hub genes for each cell type (**Figure 1B, Supplemental Figure 1**). To determine if the hub genes were prioritized for cell-type-specific functions, we assessed the area under the receiver operating characteristic curve (AUROC) score for each cell-type-specific entry. Using the Azimuth celltype database (27), which contains signature marker genes extracted from large scRNA-seq datasets, we observed a higher retrieval rate of cell-type signature genes by centrality in reference-guided CSNs than in reference-free CSNs (**Figure 1C**). Similarly, the association between the top 100 hub genes and each of the Azimuth cell-type signature genes tended to be stronger in CSNs generated via reference-guided methods than in those that relied on reference-free methods (**Supplemental Figure 2**). Among reference-guided CSNs, scHumanNet prioritized cell-type-specific genes better than SCINET, especially among B and T cells. These results suggest that scHumanNet is superior to other CSN construction methods in retrieving cell-type-specific functions of human genes.

### scHumanNet reveals commonality and differences among CSNs across cancer types

The function of tumor-infiltrating cells in cancer is often investigated using cell-type-specific gene expression. Here, we show that network biology can complement expression-based functional studies. To this end, we used scHumanNet to construct CSNs for T cells, B cells, myeloid cells, ECs, CAFs, and cancer cells from breast, colorectal, lung, and ovarian cancers. Next, we examined whether these CSNs could provide functional insights linked to cell type or disease status. Statistics for CSNs relative to each cancer type are summarized in **Supplemental Table 3**. Network comparisons across different types of non-cancerous cells revealed that only a minor portion of nodes and edges was shared across cell types in all cancers (**Figure 2A, B**); whereas a large portion was shared across cancer types (**Figure 2C, D**; **Supplemental Figures 3, 4**). These results indicate that CSNs generated by scHumanNet are shaped primarily by the cellular context rather than the disease or tissue context. Notably, the ratio of unique edges to shared edges across cancer was larger than that of unique nodes to shared nodes in all cell types, indicating that networks for the same cell type are rewired in different tissue and disease contexts.

**Figure 2.**
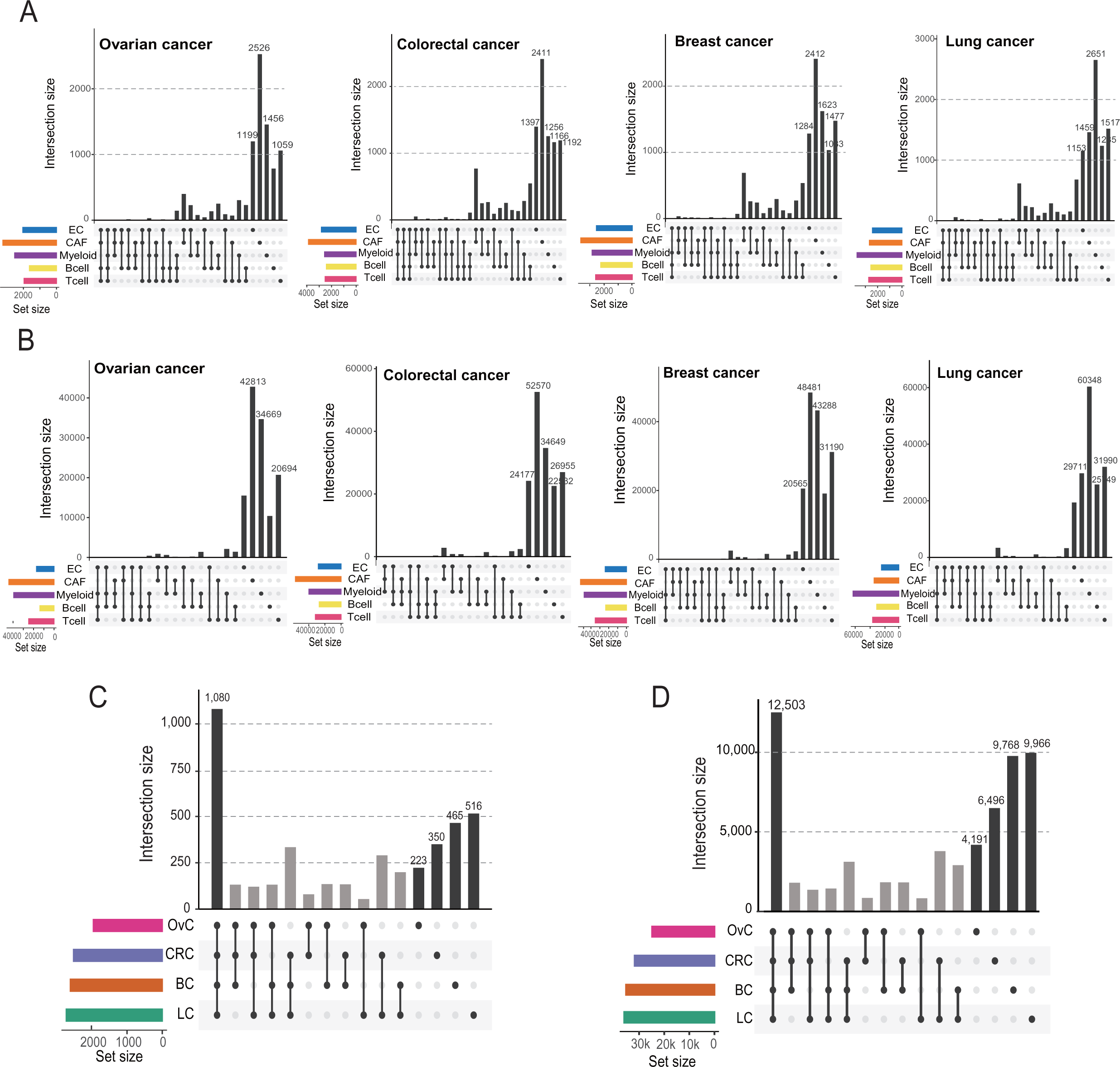
Commonality and differences between CSNs generated by scHumanNet across cancer types. **A, B**. Upset plots for five CSNs (T cells, B cells, myeloid cells, CAFs, and ECs) show overlap with respect to nodes (**A**) and edges (**B**) across cancer types. **C, D.** Node (**C**) and edge (**D**) overlap of T-cell networks among four cancer types (OvC, ovarian cancer; CRC, colorectal cancer; BC, breast cancer; LC, lung cancer).

**Figure 3.**
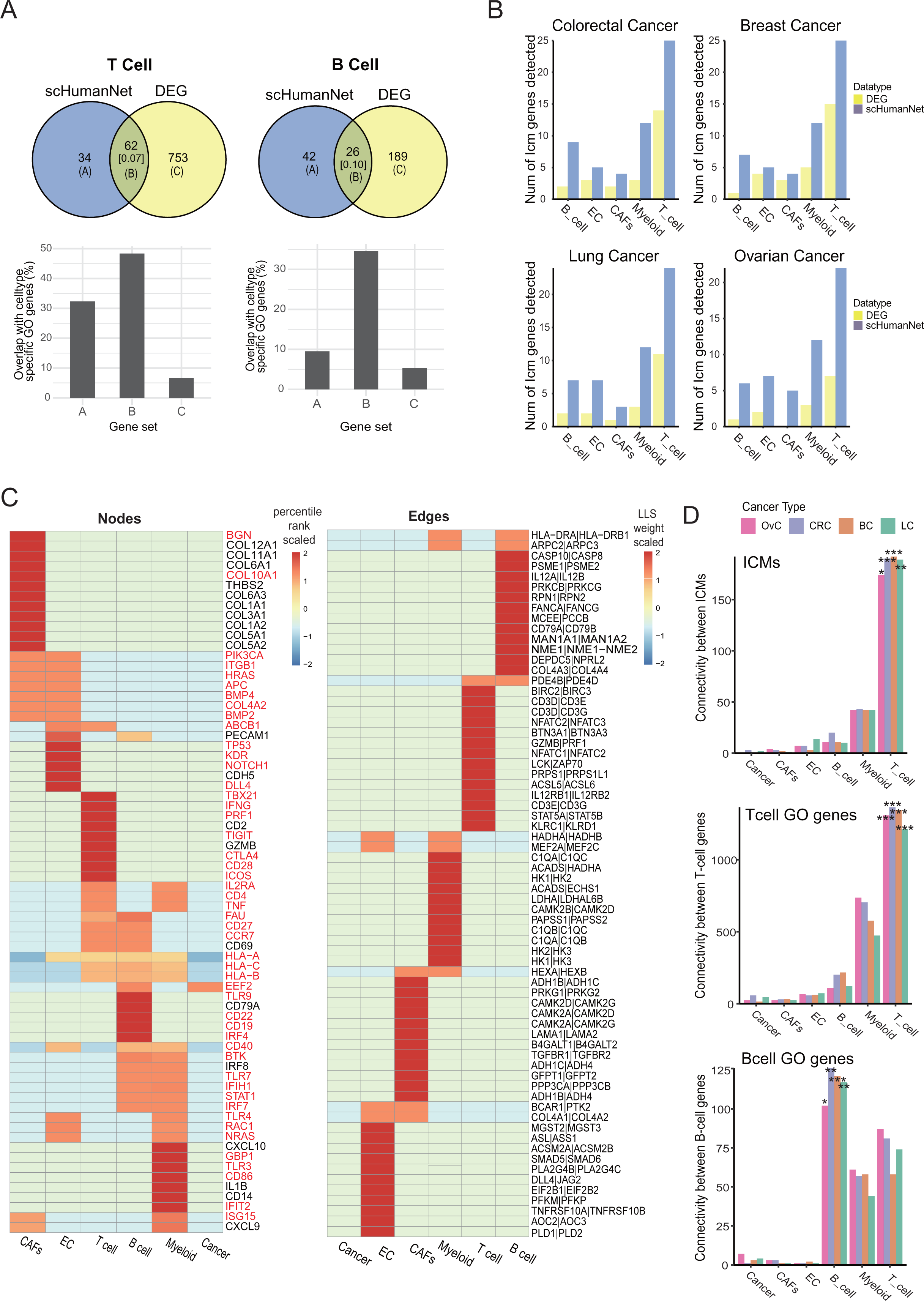
Prediction of cell-type-associated genes via differential expression analysis and network centrality by scHumanNet. **A.** Overlap of significant DEGs and significant hub genes for B- and T-cell networks in breast cancer (*q*- value < 0.05). The numbers in square brackets correspond to Jaccard indices. Overlap of genes specific for B- and T-cell functions was assessed for prediction by hub genes and DEGs (set A and set C) and their intersection (set B). **B.** Number of ICMs retrieved via *FindAllMarkers()* (DEG) and *FindDiffHubs()* (scHumanNet) in different cancer types. **C.** Heat map showing the percentile rank of top 15 hub genes (nodes) and interactions (edges) of each breast cancer network. Values were scaled per row. Results for other cancer types are reported in **Supplemental Figure 6**. Genes highlighted in red were not among the top 50 DEGs retrieved by the *FindAllMarkers()* function in the Seurat package. **D.** Within-group connectivity between ICMs, T-cell GO genes, and B-cell GO genes for all annotated cell types in scHumanNet and for each cancer type (OvC, ovarian cancer; CRC, colorectal cancer; BC, breast cancer; LC, lung cancer). \**p* < 0.05, \*\**p* < 0.01, \*\*\**p* < 0.001.

**Figure 4.**
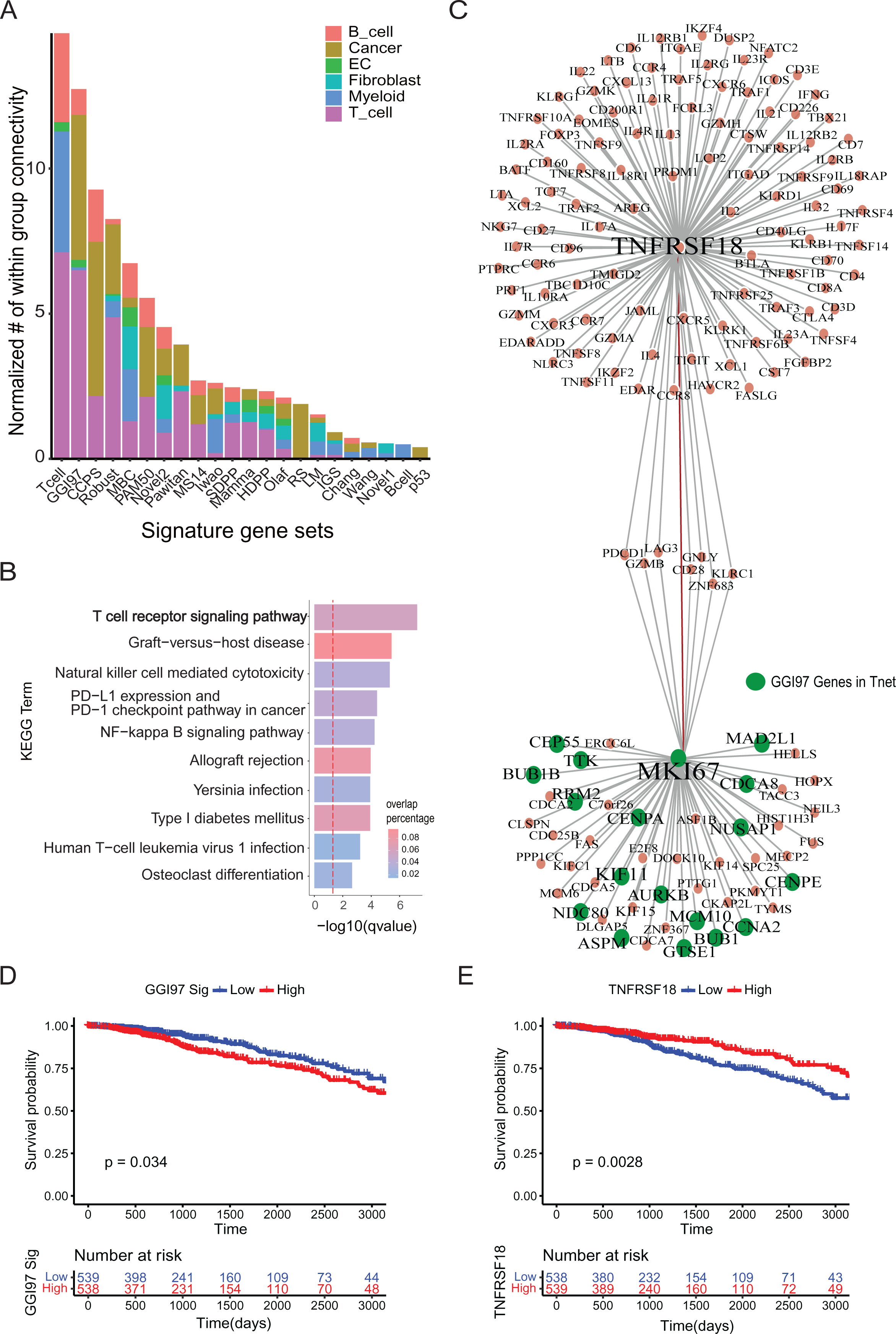
Deconvolution of breast cancer signatures to cell types with scHumanNet. **A**. Normalized within-group connectivity of each breast cancer signature in six cell-type-specific networks by scHumanNet. Within-group edge counts were normalized to the number of genes for each cell-type-specific network. Cancer signatures with at least 10 genes are presented. **B.** Gene set enrichment analysis with the KEGG pathway for the top 30 direct neighbor genes of GGI97 signature genes. The red vertical line corresponds to a *q*-value of 0.05 corrected with the Benjamini–Hochberg method. **C.** Network of genes neighboring *MKI67* and *GITR* (*TNFRSF18*) in the context of breast cancer T cells by scHumanNet. Green nodes denote GGI97 genes. **D, E.** Kaplan–Meyer plot for TCGA-BRCA cohort based on the average expression of 76 signature genes (**D**) or the expression of *GITR* (*TNFRSF18*) (**E**). Clinical samples were divided into high and low groups by median value.

### scHumanNet centrality and compactness predict cell-type specificity of gene functions

Rewiring gene interactions in different cell types might change the network centrality of genes with differential functional importance across cellular contexts. Given that hub genes with a high degree of centrality interact with many other genes in a given cellular context, we hypothesized that they were more likely to play important roles in maintaining functions specific to the given cell type. Therefore, we investigated whether scHumanNet hub genes for each type of tumor-infiltrating cell could reflect cell context-dependent functional importance across cancer types. To evaluate cell-type specificity, we utilized the GO database to collate genes reliably associated with either B or T cells (**Methods**). Next, we assessed the power of scHumanNet to predict genes specific for each cell type based on overlap with genes known to function in B or T cells. Notably, the network-based and expression-based candidate genes specific for each cell type showed low concordance (0.05 to 0.13 Jaccard similarity index), indicating complementarity of the two predictions (**Figure 3A, Supplemental Figure 5**). Moreover, the intersection between the two predictions showed strong overlap with known cell-type-specific genes. Interestingly, for the most part, network-based predictions showed a similar or higher overlap with known cell-type-specific genes than expression-based predictions, further confirming that scHumanNet hub genes could effectively identify cell-type-specific genes.

We anticipated that ICMs would be enriched among genes specific for tumor-infiltrating cells. Hence, we compiled 43 previously identified ICMs (**Supplemental Table 4**) (23) and compared their overlap with scHumanNet hubs and DEGs across cell types. For all cell types, we observed higher retrieval of ICMs by scHumanNet hub genes than by DEGs (**Figure 3B**). Notably, in all cancer types and cell types, the ICMs retrieved by DEGs were subsets of those retrieved by scHumanNet (**Supplemental Data 1**).

We prioritized genes using weighted degree of centrality in the CSNs constructed by scHumanNet and found that it was highly predictive of cell-type-specific hallmark genes (**Supplemental Data 2**). Based on this observation, we chose to more closely investigate TFs, which are key determinants for the differentiation and maintenance of particular cell identities. Cell-type-specific differential expression analysis is often insufficient to detect TFs for a given cell type because of a generally low basal level of expression. Instead, a network-based approach has been widely used to infer TF-target interactions (28, 29). We hypothesized that network centrality in CSNs could effectively prioritize TFs specific for a certain cell type. To evaluate the prediction of TFs specific for cell types by DEGs and scHumanNet centrality, we retrieved cell-type-specific TFs from TF-Marker (21), a manually curated cell-type-specific TF database. Because of the limited number of entries, we could analyze only TFs specific for B and T cells. By comparing the enrichment of known cell-type-specific TFs among the top 100 prioritized genes by scHumanNet centrality with those identified by differential expression, we found that the network-based approach was consistently better at prioritizing TFs in both B and T cells across cancer types (**Methods, Supplemental Data 3**).

We also found that scHumanNet centrality could predict cell-type-specific disease-associated genes. For example, the top 15 hub genes in T cells from all types of cancers included those involved in cell-mediated immunity (*GZMB*, *PRF1*, and *IFNG*) and immune checkpoint signaling pathways (*TIGIT* and *CTLA4*) (**Figure 3C, Supplemental Figure 6, Supplemental Data 2**). Notably, four of the five hallmark genes for T-cell immunity (*PFR1*, *IFNG*, *TIGIT*, and *CTLA4*) were not found among the top 50 DEGs (**Supplemental Data 4**). In B cells, *TLR7* and *TLR9* were found to be pan-cancer central genes but were not detected as DEGs. In myeloid cells, the top 50 pan-cancer central genes included seven genes involved in myeloid cell differentiation (*CD4*, *FCER1G*, *IRF8*, *TYROBP*, *TLR2*, *TREM2*, and *ITGAM*), but only two of them (*FCER1G* and *TYROBP*) were found among the top 50 DEGs. In CAFs from ovarian cancer, but not from other cancer types, 11 aldehyde dehydrogenase genes (*ALDH1L1*, *ALDH1L2*, *ALDH3A2*, *ALDH1A3*, *ALDH1A1*, *ALDH1A2*, *ALDH2*, *ALDH1B1*, *ALDH4A1*, *ALDH9A1*, and *ALDH6A1*) were prioritized in the top 100 hub genes by scHumanNet. Aldehyde dehydrogenase has been associated with poor survival as it promotes tumor growth in ovarian cancer (30). Notably, none of the 11 aldehyde dehydrogenase genes were among the top 100 DEGs in CAFs from ovarian cancer. The *NOTCH1* gene is expressed in ECs, where it promotes metastasis (31). We found *NOTCH1* among the top 20 hub genes in endothelial CSNs for all four cancer types (7th for breast, 13th for colorectal, 11th for lung, and 19th for ovarian cancers), but not among the top 200 DEGs in all cancer types. These results suggest that network centrality using scHumanNet can be more effective than differential expression analysis in identifying genes that play important roles in a given cellular context. These results also suggest that *FindAllHubs()* in scHumanNet can identify hub genes with cell-type-specific functions in both healthy and disease contexts.

Rewiring molecular networks across different cell types may result in differential within-group connectivity (or compactness), which can also be used to estimate functional relevance. As a proof-of-concept, we utilized ICM genes and genes specific to B and T cells. The *Connectivity()* function in scHumanNet tests the significance of within-group connectivity against a nonparametric null model using restricted random sampling that does not require the identification of optimal parameters (**Methods**). As expected, ICM genes and those specific for B and T cells were associated with T-, B-, and T-cell types, respectively, in all cancer types (**Figure 3D**). This suggests that the network-based approach provides a complementary and intuitive method for assigning gene sets to functionally relevant cell types based on compactness.

### Cell type deconvolution of cancer prognostic signatures using scHumanNet

ICMs showed the highest compactness in the T-cell network, which is consistent with their cellular role. We hypothesized that we might deconvolve disease-associated gene signatures obtained from bulk tissues into individual cell types using their network compactness across CSNs by scHumanNet. For example, cancer prognostic signatures are presumably associated with cancer cells because they are typically identified in tumor tissues. However, tumor tissues often contain also non-cancerous cells, such as stromal and immune cells, and some prognostic genes may exert their functions in non-cancerous cells of the tumor microenvironment. To test this hypothesis, we examined 33 prognostic signatures reported in breast cancer (24). We measured the normalized within-group-connectivity of each prognostic signature across the CSNs using scHumanNet (**Figure 4A**). As expected, we observed strong network compactness for many prognostic signatures from non-cancerous cells, particularly from T cells for the ‘T-cell metagene signature’ (Tcell) (32), ‘97-gene genomic grade index’ (GGI97) (33), ‘127-gene classifier’ (Robust) (34), and ‘64-gene expression signature’ (Pawitan) (35). These results indicate that T-cell function may in part account for the clinical outcomes of breast cancer.

Next, we focused on the GGI97 signature (**Supplemental Figure 7A**), which has been extensively studied and clinically validated to have an inverse correlation with survival and a positive association with chemotherapy response (36). GGI97 genes were mostly associated with the cell cycle and G2/M checkpoint pathways (47/76 genes) (**Supplemental Data 5**). Additionally, the 24 GGI97 genes detected in the T-cell network were closely connected to each other (*p* = 0.0001) (**Supplemental Figure 7B-C**) and significantly enriched in cell cycle-related functions (*p* = 0.0075 by hypergeometric test) (**Supplemental Table 1**), suggesting a role for T cell cycle control in antitumor activity. We also found that many GGI97 genes were connected to genes with a high degree of centrality and important for T cell antitumor activity (*GZMB*, *PDCD1*, *KLRC1*, *TNF*, and *ICOS*) (**Supplemental Figure 7D**). In particular, genes directly connected to GGI97 signature genes were enriched in the T cell receptor signaling pathway (**Figure 4B**), indicating that a high GGI97 score primed the immune system for a better response to chemotherapy (37).

T cell proliferation is important in the immunotherapy response. Out of 24 GGI97 signature genes in the T-cell network, 18 were direct neighbors of *Ki67* (**Figure 4C**), a known marker of cell proliferation. The GGI97 signature is associated with poor survival, which was confirmed by the median expression of GGI97 genes in TCGA-BRCA samples (**Figure 4D**). To understand the role of GGI97 genes in T cells, we examined the top 10 hub genes directly connected to GGI97 genes in the T-cell network. Notably, *GITR* (*TNSFR18*), a hub gene directly connected to *Ki67*, was prognostic of positive clinical outcomes (**Figure 4E, Methods**). Importantly, the expression of *GITR* did not correlate with the abundance of T cells (**Methods**), ensuring that we observed the cellular effect of *GITR* regardless of T-cell composition in each tumor sample (**Supplemental Figure 7E**). *GITR* has a co-stimulatory role (38) which is essential for CD8^+^ T cells to mount an antitumor immune response. When T cells bind to the ligand *GITRL*, *GITR* promotes the proliferation of effector T cells and dampens the suppressive activity of regulatory T cells (39). The GGI97 signature is predictive of chemotherapy responses. Chemotherapy can promote the cancer-immunity cycle by releasing neoantigens from dead cancer cells. Thus, the beneficial effect of *GITR* can be explained in terms of antitumor immunity. Moreover, we believe that the prognostic effect of the GGI97 signature in chemotherapy is tied to T-cell function via *GITR*. Consistent with our results, *GITRL* combined with anti-PD1 immunotherapy was shown to be effective against breast cancer, resulting in enhanced T-cell activation, proliferation, and memory differentiation (40). Taken together, our findings demonstrate that scHumanNet can deconvolve cancer prognostic signatures into cell types and identify key targets for therapeutic approaches in specific cell types.

### Identification of disease-associated cell types using differential hubness analysis in scHumanNet

Another application of scHumanNet is the identification of differential hubs, that is, genes whose centrality changes significantly between two biological contexts, such as disease and healthy conditions. The *FindDiffHub()* function in scHumanNet assigns ranks to the genes based on the degree of centrality in each context-specific network, and then identifies those genes whose percentile rank has changed significantly compared to a null model. In addition, the *TopDiffHub()* function allows users to extract the top *n* differentially ranked genes (**Methods**). Using differential hubness analysis with scHumanNet, we investigated ASD, a neurodevelopmental disorder with strong heritability (41). ASD is characterized by difficult social interaction and communication, repetitive behavior, and/or sensory susceptibility, and is likely to have many different genetic and environmental causes. A large cohort study by the SFARI consortium identified 1,231 genes (42). However, the mechanisms of action of most genes remain poorly understood. We hypothesized that, in the disease condition, perturbation of SFARI genes could result in cell-type-specific loss of wild-type molecular interactions. Thus, a decrease in network centrality could point to disease-associated cell types. Using a published dataset containing 104,559 cells from 15 donors diagnosed with ASD and 16 matching controls (11), we constructed seven CSNs for both healthy and disease conditions (**Figure 5A, Supplemental Table 5, Methods**). We found that the scHumanNet hub genes for each cell type were relevant to cell-type-specific functions (**Figure 5B**). For example, the NMDA receptor subunit *GRIN2B* is a hub gene in both excitatory and inhibitory neurons, and the TF *SOX9* is a hub gene in astrocytes (43). We also observed *CD163*, *FCER1G*, and *CD14* as hub genes in microglia (44). Interestingly, unlike the immune cell dataset, whereby a few hub genes were also detected as marker genes via DEGs, most hub genes of the brain scHumanNets were not prioritized via differential expression analysis (**Supplemental Data 6**) and, indeed, showed minimal overlap with cell-type-specific DEGs (**Supplemental Figure 8**).

**Figure 5.**
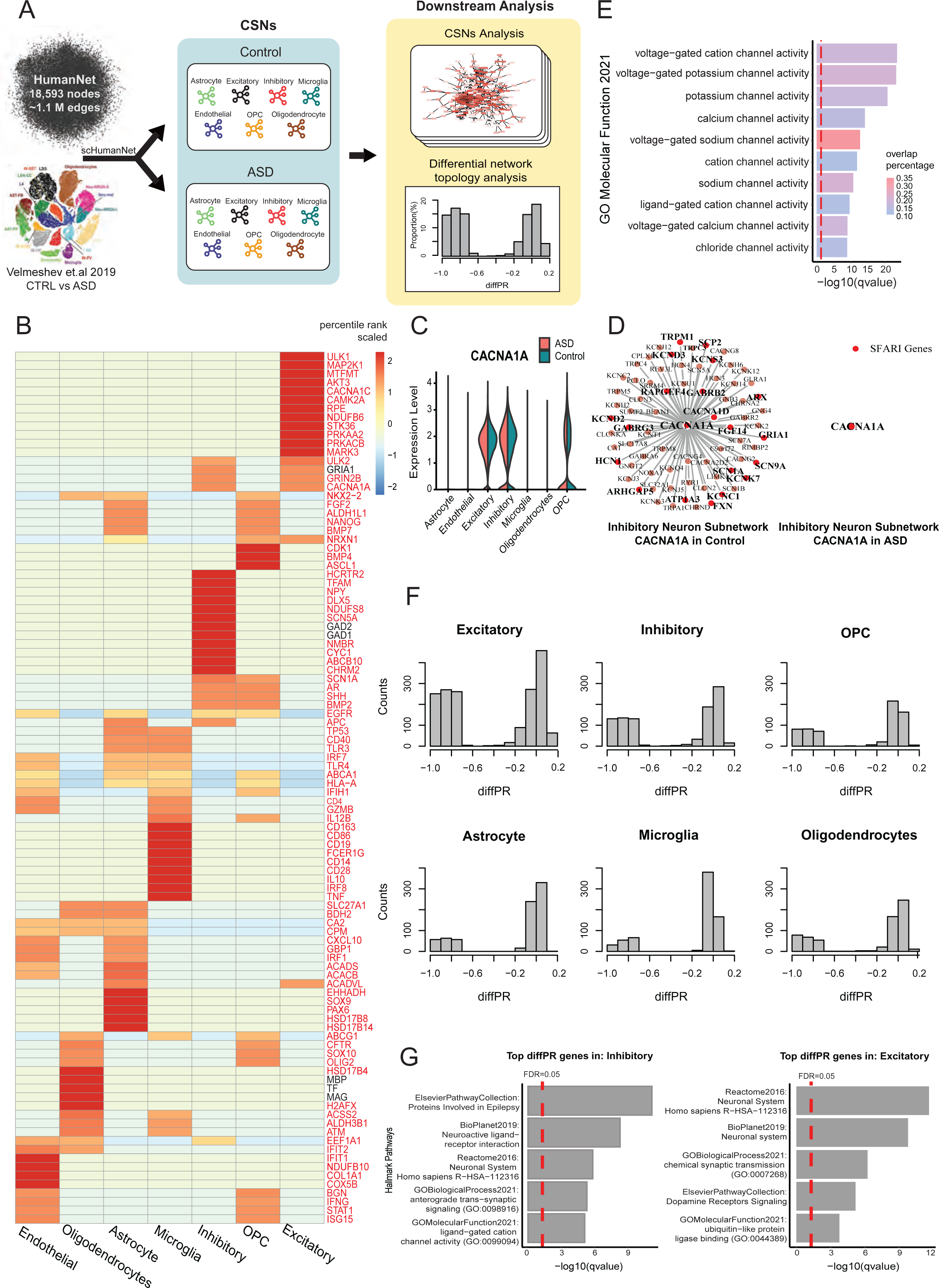
Differential hubness analysis between ASD and healthy control samples across CSNs by scHumanNet. **A**. Overview of differential hubness analysis by scHumanNet. Seven cell types were grouped and CSNs in normal and ASD conditions were constructed. **B.** Top 15 hub genes in the combined (control and ASD) networks for seven cell types. Genes highlighted in red were not among the top 50 DEGs identified by the *FindMarkers()* function in the Seurat package. **C.** Violin plot showing the normalized expression of *CACNA1A* for each cell type in ASD and healthy conditions. The statistical significance of differences between cell types was not evaluated. **D.** Network visualization of *CACNA1A* and neighboring genes in healthy (left) and ASD (right) inhibitory neurons by scHumanNet. SFARI genes are in red (20 genes out of 72 neighbors in the healthy control, none in ASD). **E.** Direct neighbors of *CACNA1A* from normal inhibitory neurons by scHumanNet were assessed for enrichment using the GOBP database. The red vertical line corresponds to a *q*-value of 0.05 corrected with the Benjamini–Hochberg method. **F.** Distribution of diffPR values for genes with hubness (PR) > 0.7 in control cell types. **G.** Hallmark pathways of genes in ASD derived from five pathway databases (Reactome, BioPlanet, Elsevier Pathway Collection, GO Biological Process, GO Molecular Function) and identified in inhibitory neurons (left) and excitatory neurons (right).

Our analysis also revealed that many genes differed significantly in terms of network centrality between the control and disease conditions, despite modest fold changes (**Supplemental Figure 9A**). By assessing genes with the highest differences in centrality rank via *FindDiffHub()* with default parameters (**Methods**), we found that differential hubs from excitatory and inhibitory neurons were significantly enriched with SFARI genes, which contrasted with DEGs being found mostly in ECs and astrocytes (**Supplemental Data 7, 8**). In particular, the highest overlap between differential hub genes and SFARI genes was observed in excitatory neurons (**Supplemental Data 8**), although several key ASD genes, including *GRIN2B* and *MECP2* (45, 46), were found as differential hubs in inhibitory neurons. Even though *GRIN2B* and *MECP2* are expressed in both excitatory and inhibitory neurons, they were found to be differential hubs only in the latter (**Supplemental Figure 9B**), implying that they may be functionally more important in inhibitory neurons. This finding has been experimentally validated in a mouse model (47) and suggested by a human study (48), in which inhibitory neurons were enriched for overexpressed SFARI genes. Similarly, for *CACNA1A*, we found that although it was not differentially expressed in inhibitory neurons (**Figure 5C**), there was a significant difference in terms of network centrality (**Figure 5D**), and many of the functional interactions were lost in the ASD inhibitory neuron network. The interacting genes were mostly associated with ion channels (**Figure 5E**), suggesting that the function of neural regulation, especially in inhibitory neurons, might be impaired by*CACNA1A* loss-of-function mutations (49). These results demonstrated that differential hubness analysis using scHumanNet could reveal disease-associated cell types.

Finally, we investigated whether genes with high centrality in healthy conditions but low centrality in disease conditions might provide insights regarding cell-type-specific disease mechanisms (**Methods**). We found that excitatory neurons, inhibitory neurons, and oligodendrocyte progenitor cells had the highest frequency of loss-of-function genes compared to other cell types (**Figure 5F**). Notably, genes with high centrality in disease but low centrality in healthy controls were less frequent across all cell types (**Supplemental Figure 10A**). Gene set enrichment analysis of hubs lost in neurons revealed that their function was primarily associated with neuronal activity (**Figure 5G**). For inhibitory neurons, the hub genes lost under healthy conditions were enriched in ‘increased anxiety-related response’ (MGI Phenotype), ‘anterograde trans-synaptic signaling’ (GOBP), and ‘ligand-gated cation channel activity’ (GOMF). In excitatory neurons, the genes that lost centrality were enriched in ‘chemical synaptic transmission’ (GOBP), ‘dopamine receptors signaling’ (Elsevier Pathway Collection), and ‘protein secretion’ (MSigDB Hallmark). These results imply that, in disease conditions, these hub genes lost most of their interactions with other genes, resulting in the dysregulation of neuronal function in ASD. In contrast, genes that became more central in ASD networks were not enriched in pathways related to neuronal function (**Supplemental Figure 10B**).

## Discussion

The main goal of single-cell biology is resolving the cellular heterogeneity of human diseases. Single-cell gene expression analysis may enable the identification of disease-associated cell types based on the differential expression of disease-associated genes in specific cell types. In the present study, we described scHumanNet, a computational platform for network-based analysis of cell-type specificity, which can complement expression-based approaches. The core component of this platform is the reconstruction of CSNs, gene network specific to distinct cell types. Single-cell transcriptome data have been utilized to construct CSNs with either reference-guided or reference-free network inference methods. The evaluation of inferred CSNs is not a trivial task because of the lack of high-quality and experimentally validated gene-gene interactions for particular cell types. In fact, because of the high false positive rate of inferred gene-gene interactions from single-cell transcriptome data, functional hypotheses from these networks are generally based on a group of edges rather than individual ones. Here, we validated the quality of CSNs by the retrieval of cell-type-specific genes among hub genes and network compactness of functional genes in the corresponding cell types. In the present study, we compared various approaches for CSN inference from single-cell transcriptome data and found that reference-guided methods outperformed reference-free methods. These results can be explained by the noisy and sparse nature of single-cell transcriptome data, which generate many false-positive gene-gene interactions (4). Furthermore, among the two reference-guided CSN analysis platforms, scHumanNet was superior to SCINET. Although they utilized the same network inference algorithm, they employed different reference interactomes. Previously, we demonstrated that HumanNet, the reference interactome of scHumanNet, performed significantly better than other human gene networks, including the reference interactome of SCINET, in predicting disease genes (8). This indicates that the quality of the reference interactome is key to the performance of reference-guided CSNs, and future improvement of the former will further ameliorate CSNs.

In this study, we have demonstrated two applications of CSNs in the investigation of cell-type specificity of human disease genes. First, the effects of disease genes can be deconvolved into cell types based on the network compactness of a group of disease genes across CSNs. For example, cell-type deconvolution of breast cancer prognostic signatures showed high compactness not only in cancer cells but also in other tumor-infiltrating cells such as immune cells. The importance of T cells in antitumor activity may account for the large functional bias of prognostic genes towards T cells. Indeed, one of the identified hub genes was *GITR*, a T-cell-specific regulator that plays an important role in the survival of patients with breast cancer. We believe that our network-based approach for associating gene sets with cell types can complement expression-based methods, such as GSVA (50) and scfind (51). In the future, we may expand the scHumanNet platform to systematic cell type deconvolution of disease gene sets for all cell types of each tissue and thus generate CSNs for human cell atlas data. Second, we utilized CSNs to identify disease-associated cell types based on differential hubness between disease and healthy conditions across cell types. Therefore, the scHumanNet platform allows the analysis of differential hub genes. Using the scHumanNet pipeline, we identified inhibitory neurons as a major cell type associated with ASD. These results suggest that a network-based approach can complement an expression-based approach to identify disease-associated cell types using single-cell transcriptome data.

There are some limitations to scHumanNet. Although our results suggest that the reference-guided method yields more biologically relevant CSNs, it comes at the expense of being unable to discover novel interactions specific for the cell type. In addition, cell type deconvolution may be unreliable with a small group of genes (e.g., a set of three genes) because a statistical test for network compactness requires a relatively large number of genes to ensure a sufficient degree of confidence. Further studies are required to address these shortcomings.

In conclusion, we present scHumanNet, a computational platform for single-cell network biology, capable of resolving the cellular heterogeneity of disease-related gene functions. We demonstrate that scHumanNet can deconvolve the functional effect of disease gene sets into cell types and identify disease-associated cell types via topological analysis of CSNs. These results suggest that scHumanNet will boost our understanding of cell-type specificity of human disease genes and thus advance precision medicine.

### Code availability

The code for scHumanNet and the codes used to generate the figures in this manuscript can be downloaded from https://github.com/netbiolab/scHumanNet.

## Acknowledgement

We thank Dr. Mohammadi for guidance in implementing the SCINET algorithm to scHumanNet, as well as Dr. Karen Dixon and Dr. Susanna Mierau for their helpful discussions and comments on the manuscript.

## Author contributions

JC and IL conceived the project. JC wrote the code and analyzed the data. JY and JWC assisted with the analysis. IL and MH supervised the research. JC, MH, and IL wrote the manuscript with input from the other authors.

---

**Supplemental Figure 1.**
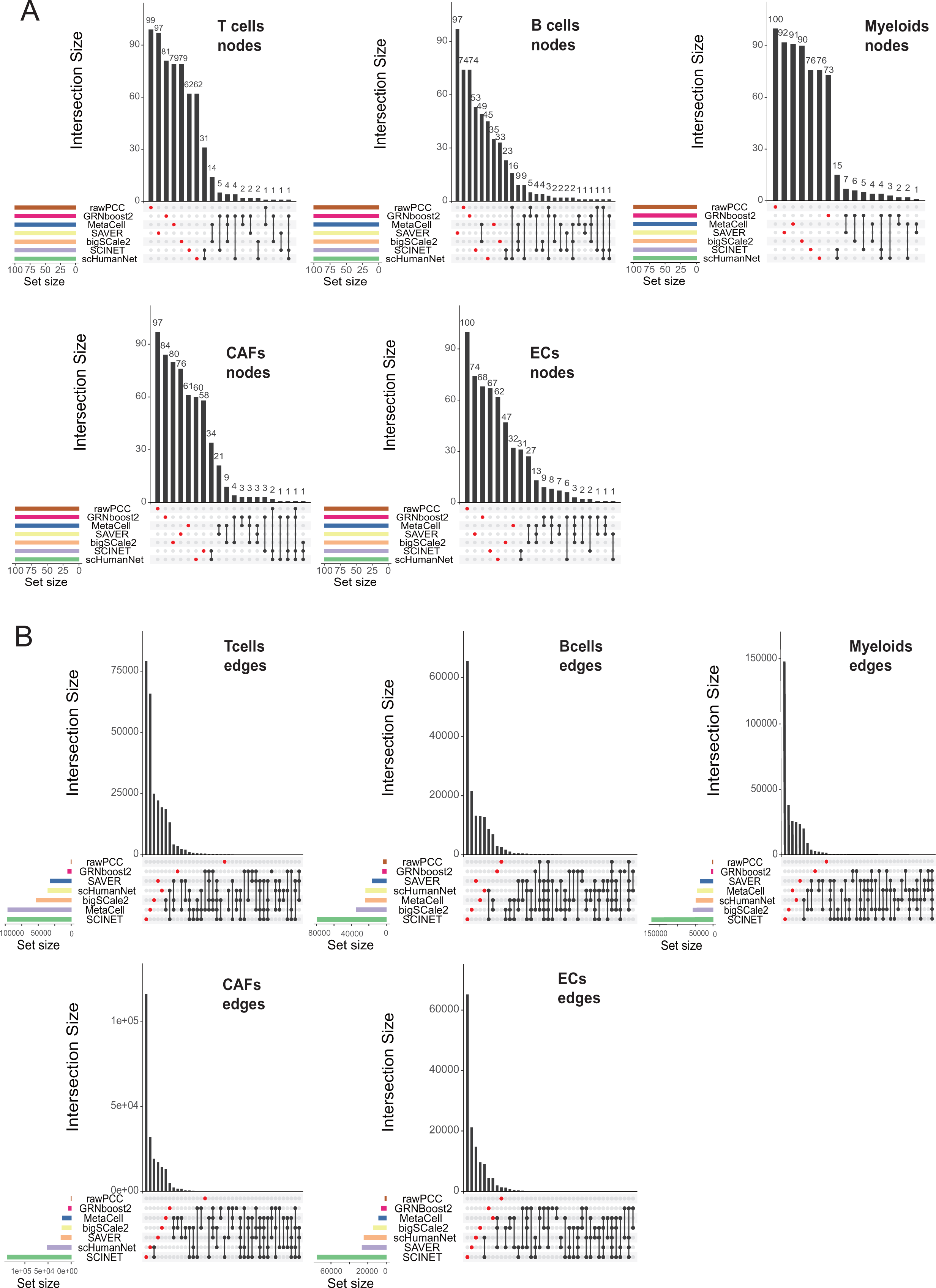
Upset plots for node and edge overlap between cell-type-specific networks (CSNs) by different methods. **A**. Upset plots for node overlap. **B.** Upset plots for edge overlap. Five cell types, including B cells, T cells, myeloid cells, cancer-associated fibroblasts (CAFs), and endothelial cells (ECs), were assessed.

**Supplemental Figure 2.**
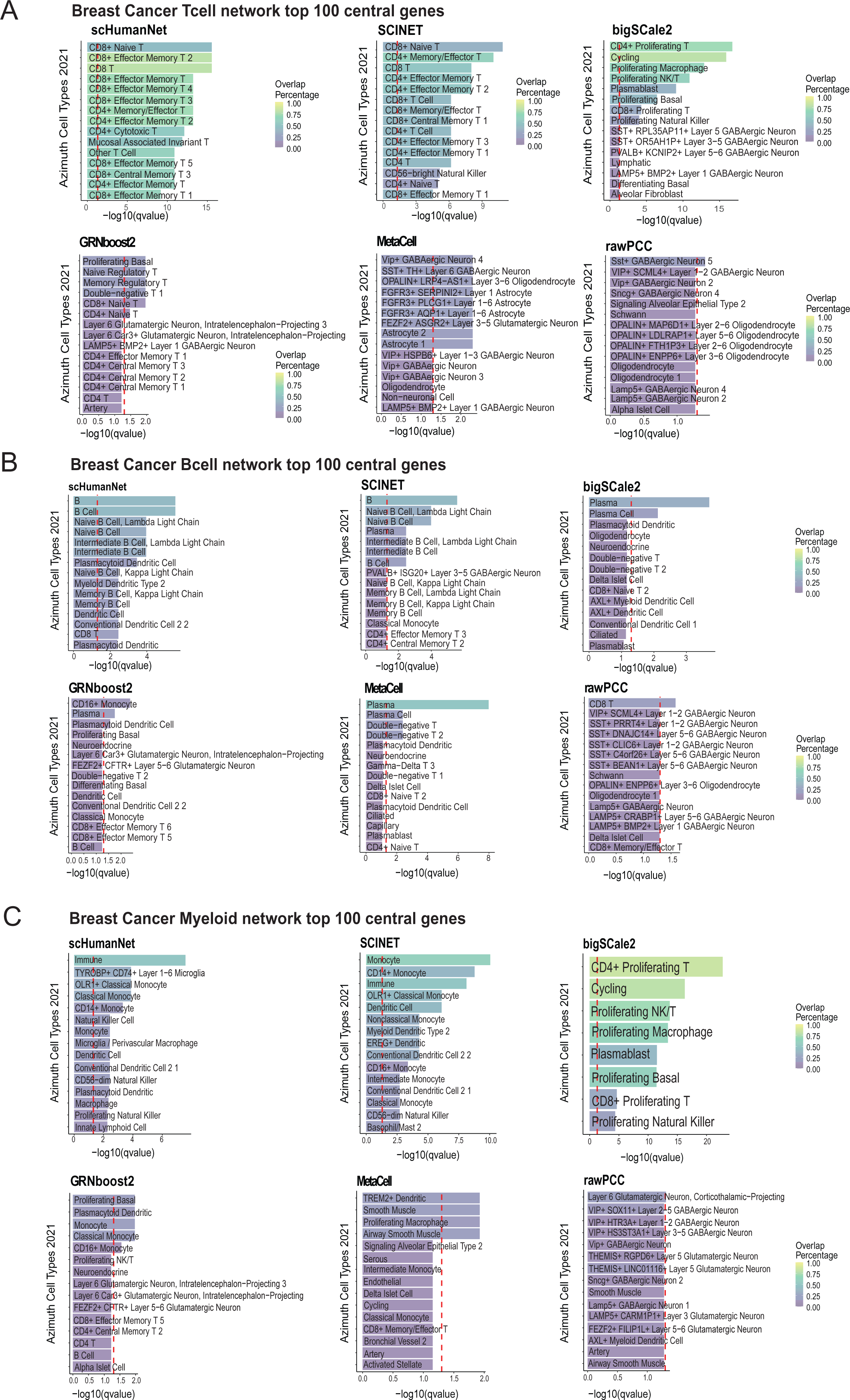
Identification of cell-type-specific genes by centrality in CSNs using scHumanNet and five other single-cell network inference methods. **A–C**. The top 100 hub genes in the networks specific for T cells (**A**), B cells (**B**), and myeloid cells (**C**) were tested for enrichment of cell-type-specific genes derived from the Azimuth celltype database. The results for networks obtained with SAVER imputation are not shown, as hub genes produced no cell-type-specific terms enriched for any cell type. The red vertical line corresponds to a *q*-value of 0.05 corrected with the Benjamini–Hochberg method.

**Supplemental Figure 3.**
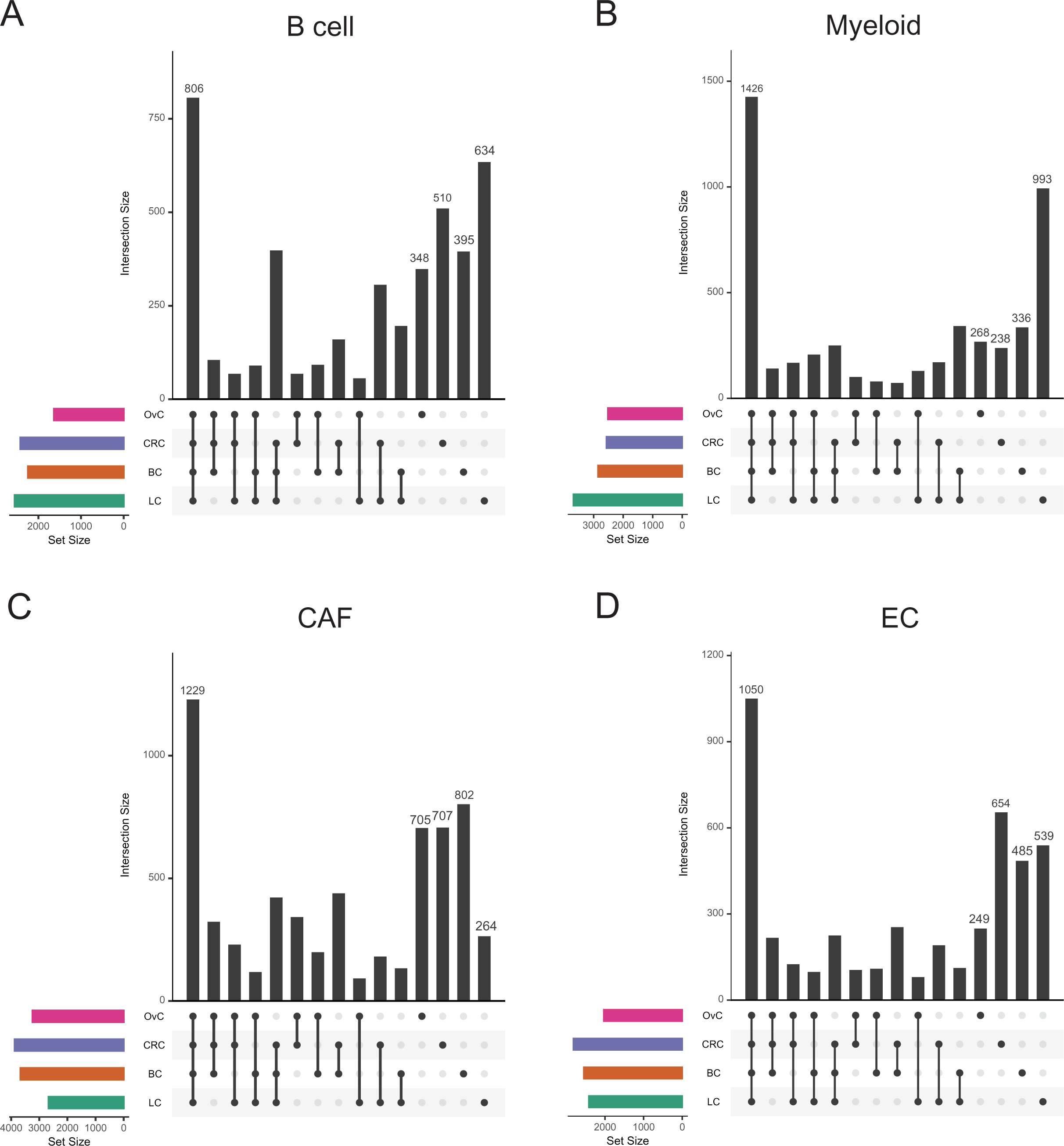
Overlap of CSN nodes by scHumanNet among cancer types. **A–D**. Upset plots for four cell types, including B cells (**A**), myeloid cells (**B**), CAFs (**C**), and ECs (**D**), showing overlap between CSN nodes among cancer types.

**Supplemental Figure 4.**
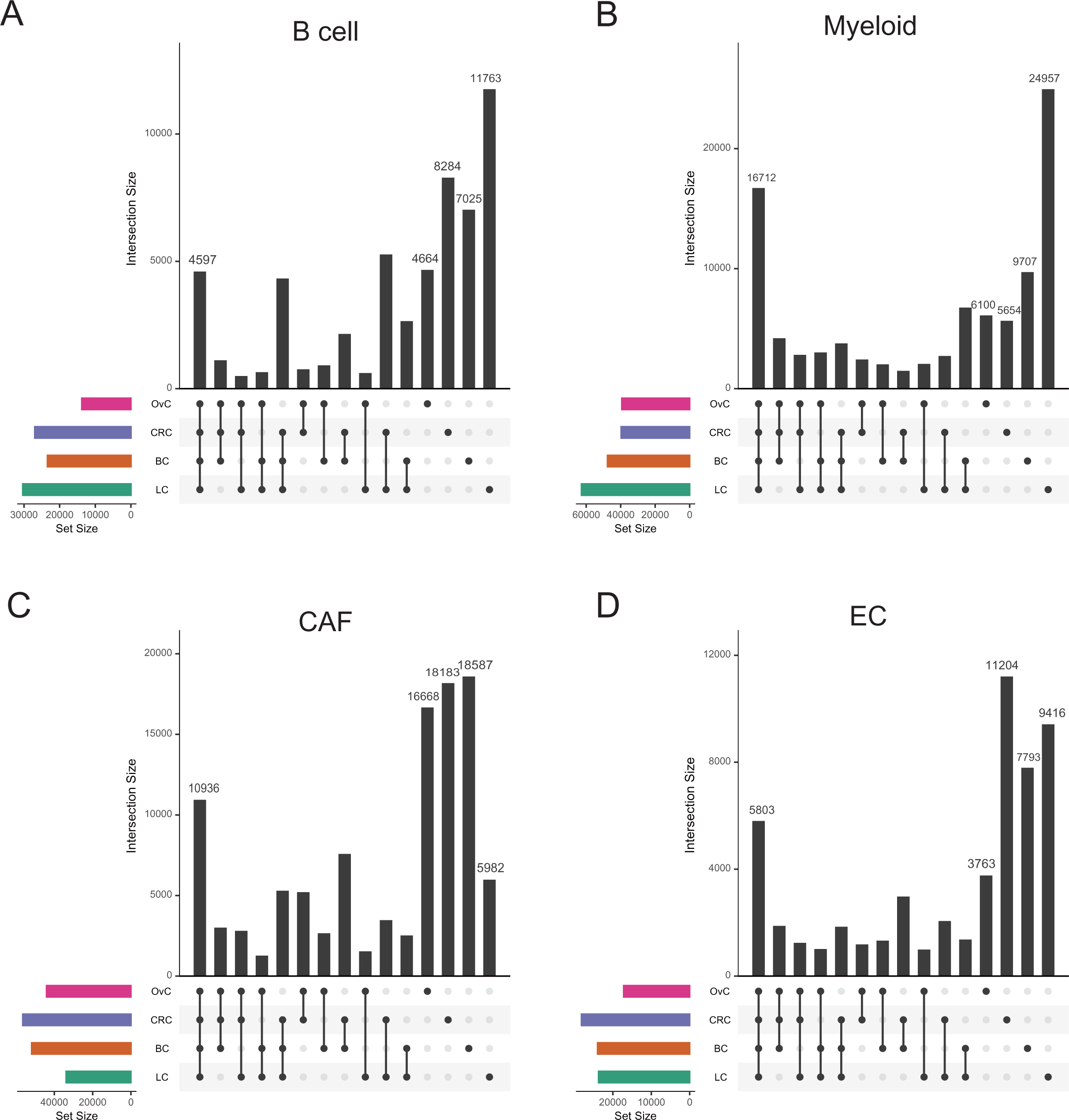
Overlap of CSN edges by scHumanNet among cancer types. **A–D**. Upset plots for four cell types, including B cells (**A**), myeloid cells (**B**), CAFs (**C**), and ECs (**D**) showing overlap between CSN edges among cancer types.

**Supplemental Figure 5.**
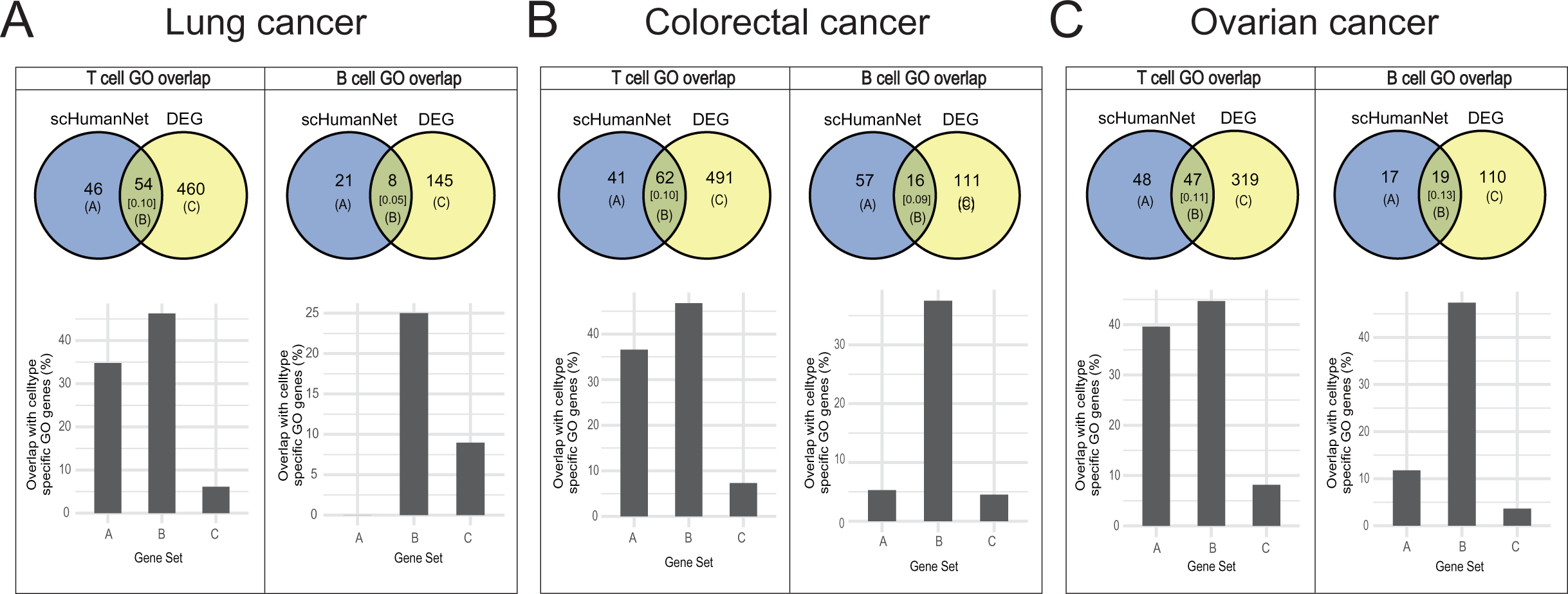
Overlap of cell-type-specific genes predicted using gene expression and network centrality. **A–C**. Venn diagram of T- or B-cell-specific genes predicted by significant DEGs and hubs in CSNs by scHumanNet for lung cancer (**A**), colorectal cancer (**B**), and ovarian cancer (**C**). The numbers in square brackets correspond to Jaccard indices. Overlap of genes specific for T- and B-cell functions was assessed for network and DEG-specific gene sets (set A and set C) and the intersection of both (set B).

**Supplemental Figure 6.**
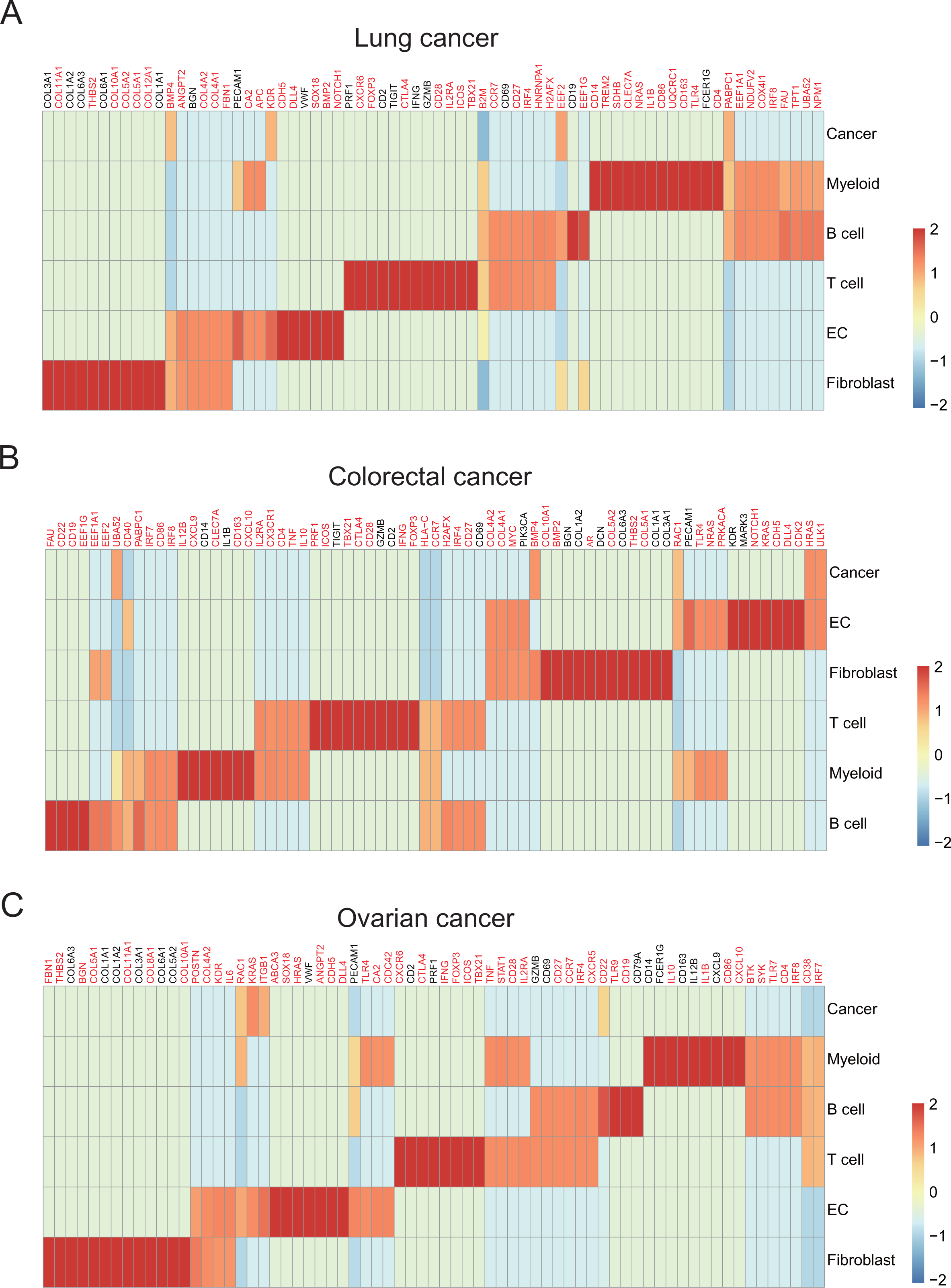
Top 15 hub genes of the CSN generated by scHumanNet for cancers. **A–C**. The top 15 hub genes were calculated as percentile ranks and scaled for lung cancer (**A**), colorectal cancer (**B**), and ovarian cancer (**C**). Genes highlighted in red were not included within the top 50 DEGs by Seurat’s *FindMarkers()* function.

**Supplemental Figure 7.**
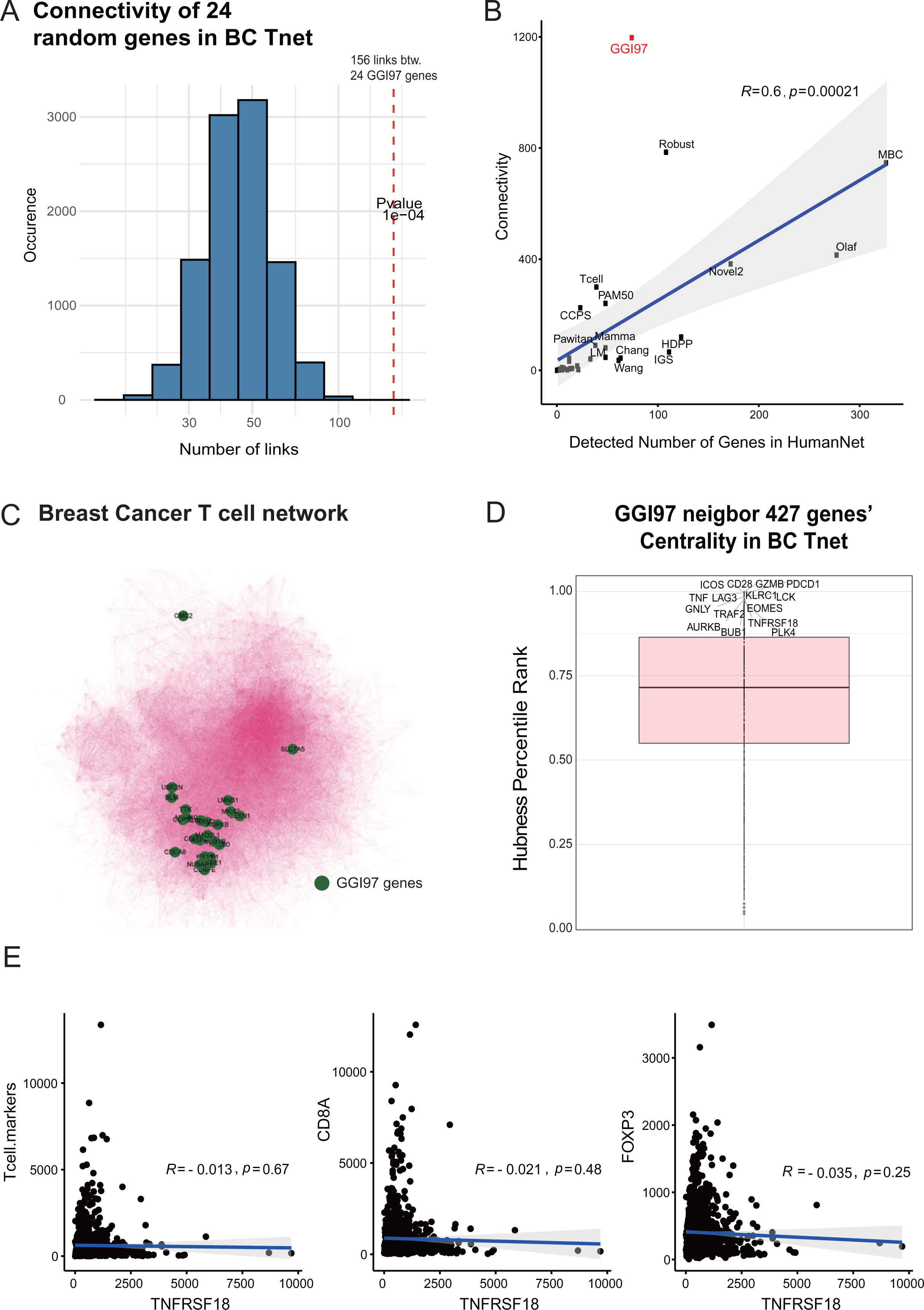
Deconvolution of breast cancer signatures into cell types by scHumanNet. **A**. Functional connectivity of 24 GGI genes detected in the T cell network for breast cancer by scHumanNet compared with the connectivity of 24 randomly selected genes. The *p*-value was calculated non-parametrically. **B.** Positive correlation between the number of signature genes detected in T-cell networks by scHumanNet and the number of connectivities. Within-group connectivity for each cell-type-specific network was normalized to network size (see Figure 4A). GGI97 showed high connectivity, despite only a moderate number of genes being detected (highlighted in red). **C.** Visualization of the entire breast cancer T-cell network with 2,611 nodes and 35,210 edges. GGI genes are highlighted in green. **D.** Percentile rank of first degree neighbors for all GGI97 genes. The top 15 genes were labeled. **E.** Correlation between TCGA-BRCA dataset with *TNFRSF18* (*GITR*) and T-cell-related signatures. TCGA-BRCA dataset was filtered for female samples and normalized using DESeq2. T-cell markers on the far right correspond to the mean expression of *CD3D, CD3E*, and *CD3G*. Pearson correlation coefficient (R) and Spearman correlation coefficient (ρ) were calculated.

**Supplemental Figure 8.**
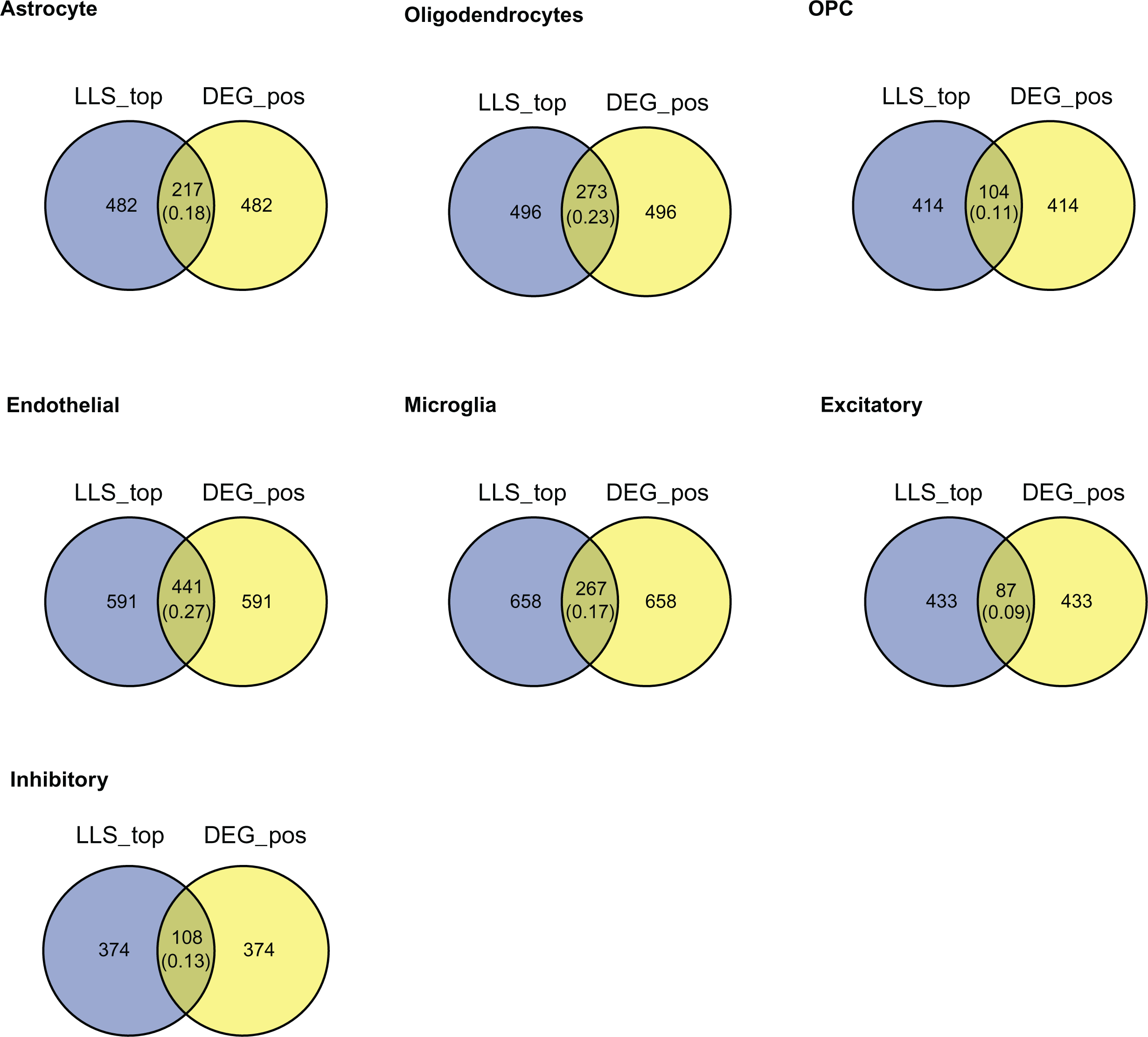
Comparison of genes prioritized by DEGs and CSN hub genes in neuronal cell types. For each cell type reported by Velmeshev *et al*. (2019), DEGs (Wilcoxon, FDR < 0.05, log fold change > 0.25) and hub genes in the CSN generated by scHumanNets were compared. The numbers in parenthesis correspond to Jaccard indices. OPC, oligodendrocyte progenitor cell.

**Supplemental Figure 9.**
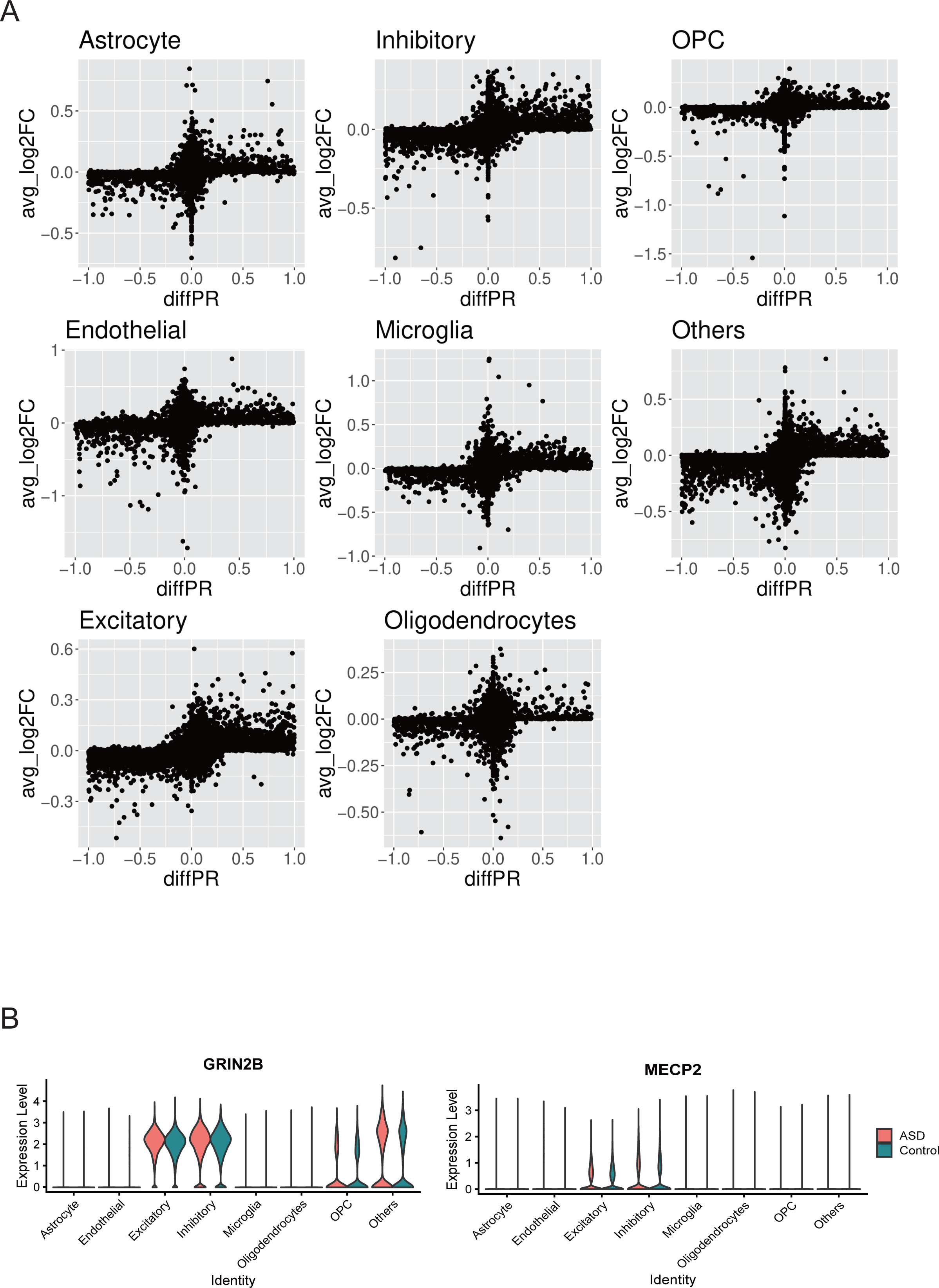
Differential hubness analysis for CSNs between control and autism spectrum disorder (ASD) conditions using scHumanNet. **A**. Evaluation of differential hubness for each gene derived from network analysis and log fold change derived from scRNA-seq expression data. **B.** Expression levels of *GRIN2B* and *MECP2* of ASD and healthy control samples are presented for each brain cell type.

**Supplemental Figure 10.**
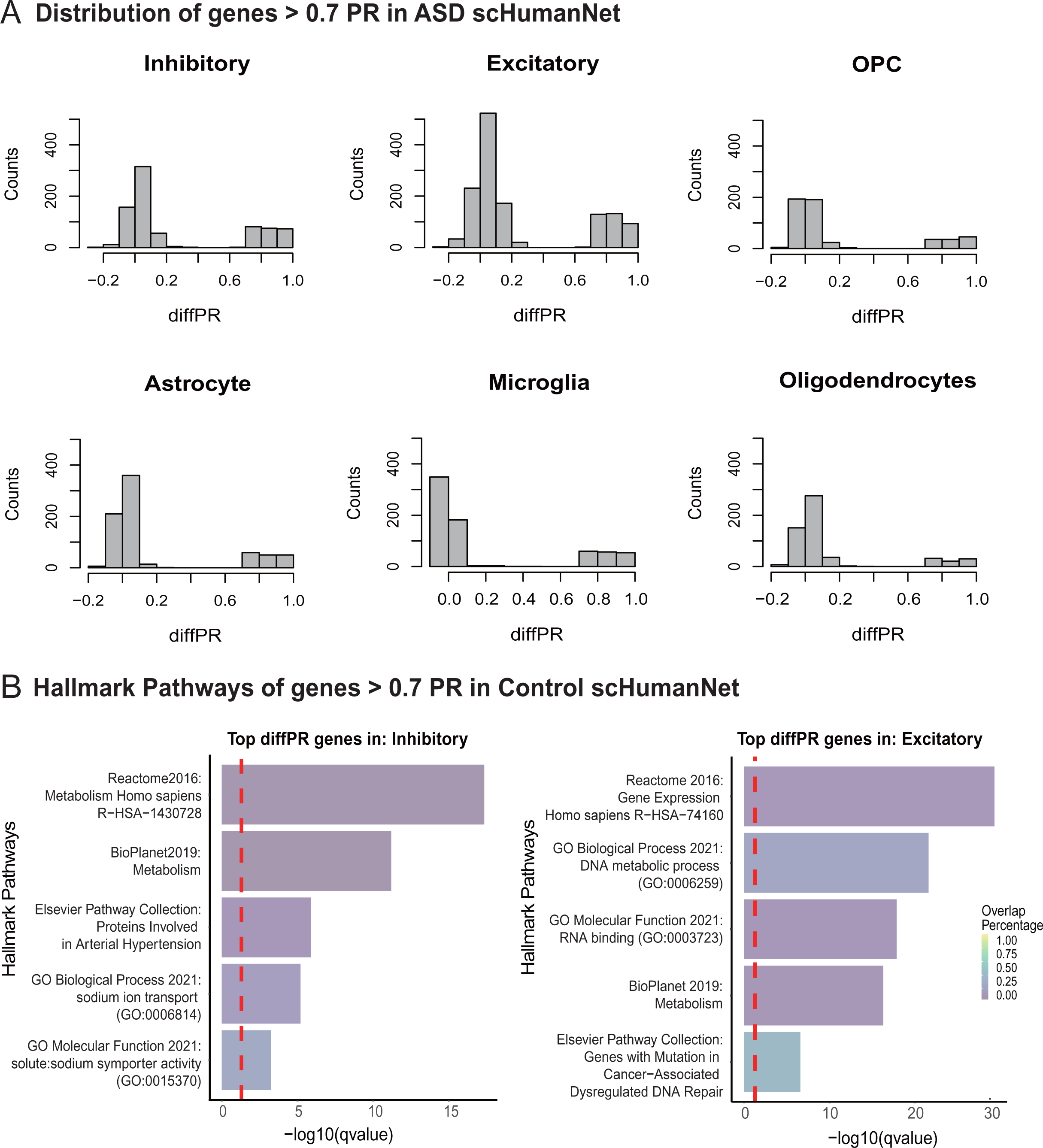
Analysis of differential hubness genes between ASD and healthy controls. **A**. Distribution of genes with > 0.7 centrality and > 0.7 differential hubness in ASD. **B.** Hallmark pathways associated with genes that have high centrality in ASD but low centrality in healthy controls based on five pathway databases (Reactome, BioPlanet, Elsevier Pathway Collection, GO Biological Process, GO Molecular Function). Pathways detected in inhibitory neurons (left) and excitatory neurons (right) are shown. The red vertical line corresponds to a *q*-value of 0.05 corrected with the Benjamini–Hochberg method.

**Supplemental Table 1.** List of genome grade index GGI97 signature genes.

|  |  |  |  |
| --- | --- | --- | --- |
| CMC2 | TTK | NUSAP1 | GTSE1 |
| ASPM | MKI67 | AURKB | RRM2 |
| SLC7A5 | KIF11 | CENPE | MCM10 |
| CEP55 | CDCA8 | NDC80 | UBE2N |
| SESN1 | CCNA2 | BUB1B | MAD2L1 |
| CENPA | BLM | LMNB1 | BUB1 |
| RACGAP1 | BIRC5 | CCNB1 | CCNB2 |
| CCNE2 | CDC2 | CDC20 | CDC25A |
| KPNA2 | MCM2 | UBE2S | MYBL2 |
| 13CDNA73 | BBS1 | BM039 | BRRN1 |
| C20orf24 | CCT5 | CDCA3 | CDK2 |
| CDKN3 | CENPF | CX3CR1 | CYBRD1 |
| DDX39 | DKFZp762E1312 | DLG7 | DONSON |
| ESPL1 | EXO1 | FEN1 | FLJ10156 |
| FLJ20477 | FLJ20641 | FLJ21062 | FLJ21827 |
| FLJ23554 | FOXM1 | GMPS | H2AFZ |
| HMGB3 | HMMR | HSMPP8 | KIF20A |
| KIF2C | KIF4A | KNSL7 | LAMB2 |
| MARS | MELK | MLF1IP | NUDT1 |
| OIP5 | ORMDL2 | PLK1 | POLQ |
| PRC1 | RNASEH2A | SHMT2 | SIRT3 |
| SPAG5 | STARD13 | STK6 | TIMELESS |
| TPX2 | TRIP13 | TROAP | TTC10 |
| ZWINT | MCM4 |  |  |

Green: detected in Breast cancer T-cell CSN
Red: either deprecated, replaced or withdrawn from the NCBI database.

**Supplemental Table 2.** Number of nodes and edges in the CSNs used in this study.

|  | # Nodes |  |  |  |  | # Edges |  |  |  |  |
| --- | --- | --- | --- | --- | --- | --- | --- | --- | --- | --- |
|  | T cells | B cells | Myeloid cells | CAFs | ECs | T cells | B cells | Myeloid cells | CAFs | ECs |
| **scHumanNet** | 2,448 | 2,066 | 2,738 | 3,504 | 2,401 | 35,120 | 23,430 | 47,678 | 51,851 | 23,994 |
| **bigScale2** | 1,890 | 1,195 | 3,928 | 1,872 | 1,757 | 53,000 | 34,000 | 56,000 | 20,000 | 14,000 |
| **SAVER** | 1,115 | 797 | 2,172 | 1,009 | 1,283 | 32,000 | 16,000 | 35,000 | 22,000 | 26,000 |
| **GRNboost2** | 2,332 | 1,840 | 2,143 | 2,795 | 2,469 | 5,302 | 3,946 | 4,539 | 6,071 | 5,354 |
| **MetaCell** | 2,739 | 1,819 | 4,133 | 1,915 | 1,103 | 96,000 | 24,000 | 44,000 | 19,000 | 8,000 |
| **rawPCC** | 121 | 716 | 530 | 134 | 557 | 221 | 3,230 | 1366 | 167 | 1313 |

CAFs, cancer-associated fibroblasts; ECs, endothelial cells.

**Supplemental Table 3.** Number of genes and edges in CSNs inferred by scHumanNet for four cancer types.

| **Cancer type** | **Cell type** | **# Genes** | **# Edges** |
| --- | --- | --- | --- |
| **Colorectal cancer** | T cell | 2,521 | 31,768 |
|  | B cell | 2,421 | 26,997 |
|  | Myeloid | 2,568 | 39,794 |
|  | Endothelial | 2,821 | 28,197 |
|  | Fibroblast | 3,873 | 56,487 |
|  | Enteric glia | 1,951 | 13,164 |
|  | Epithelial | 3,204 | 61,057 |
|  | Mast | 1,539 | 9,585 |
|  | Tumor | 4,423 | 89,859 |
| **Breast cancer** | T cell | 2,611 | 35,210 |
|  | B cell | 2,242 | 23,430 |
|  | Myeloid | 2,855 | 47,678 |
|  | Endothelial | 2,550 | 23,994 |
|  | Fibroblast | 3,665 | 51,851 |
|  | Dendritic cell | 1,996 | 24,236 |
|  | Mast | 1,522 | 8,882 |
|  | Tumor | 4,426 | 79,920 |
| **Lung cancer** | T cell | 2,726 | 35,933 |
|  | B cell | 2,554 | 30,365 |
|  | Myeloid | 3,687 | 62,810 |
|  | Endothelial | 2,420 | 23,740 |
|  | Fibroblast | 2,669 | 33,822 |
|  | Alveolar | 2,839 | 31,910 |
|  | Epithelial | 2,482 | 23,997 |
|  | Mast | 1,862 | 13,053 |
|  | Tumor | 5,075 | 111,043 |
| **Ovarian cancer** | T cell | 1,957 | 24,824 |
|  | B cell | 1,633 | 13,806 |
|  | Myeloid | 2,521 | 39,375 |
|  | Endothelial | 2,033 | 17,204 |
|  | Fibroblast | 3,238 | 44,083 |
|  | Tumor | 3,977 | 72,159 |

**Supplemental Table 4.** List of the 43 immune checkpoint molecule genes used in this study.

|  |  |  |  |
| --- | --- | --- | --- |
| ADORA2A | BTLA | CD200 | CD200R1 |
| CD244 | CD27 | CD274 | CD276 |
| CD28 | CD40 | CD40LG | CD80 |
| CD86 | CEACAM1 | CTLA4 | HAVCR1 |
| HAVCR2 | ICOS | ICOSLG | IDO1 |
| IL2RB | KIR3DL1 | LAG3 | LAIR1 |
| LGALS3 | NECTIN2 | PDCD1 | PDCD1LG2 |
| PVR | SLAMF1 | TIGIT | TNFRSF12A |
| TNFRSF14 | TNFRSF18 | TNFRSF25 | TNFRSF4 |
| TNFRSF9 | TNFSF14 | TNFSF18 | TNFSF4 |
| TNFSF9 | VSIR | VTCN1 |  |

**Supplemental Table 5.** Number of genes and links in CSNs inferred by scHumanNet for autism spectrum disorder and healthy controls.

| Condition | Cell type | # Genes | # Links |
| --- | --- | --- | --- |
| **Autism spectrum disorder** | Astrocyte | 2,505 | 26,610 |
|  | Excitatory | 4,488 | 68,448 |
|  | Inhibitory | 2,597 | 23,992 |
|  | Microglia | 2,387 | 32,270 |
|  | Endothelial | 2,394 | 35,789 |
|  | OPC | 1,784 | 14,017 |
|  | Oligodendrocyte | 1,860 | 11,629 |
|  | Others | 2,532 | 27,807 |
| **Control** | Astrocyte | 2,549 | 27,810 |
|  | Excitatory | 5,731 | 117,415 |
|  | Inhibitory | 3,164 | 36,348 |
|  | Microglia | 2,375 | 31,190 |
|  | Endothelial | 2,626 | 40,769 |
|  | OPC | 2,154 | 18,613 |
|  | Oligodendrocyte | 2,203 | 16,660 |
|  | Others | 3,234 | 50,535 |

OPC, oligodendrocyte progenitor cell.

